# Dynamic molecular mechanism of the nuclear pore complex permeability barrier

**DOI:** 10.1101/2023.03.31.535055

**Authors:** Toshiya Kozai, Javier Fernandez-Martinez, Trevor van Eeuwen, Paola Gallardo, Larisa E. Kapinos, Adam Mazur, Wenzhu Zhang, Jeremy Tempkin, Radhakrishnan Panatala, Maria Delgado-Izquierdo, Barak Raveh, Andrej Sali, Brian T. Chait, Liesbeth M. Veenhoff, Michael P. Rout, Roderick Y. H. Lim

## Abstract

Nuclear pore complexes (NPCs) mediate nucleocytoplasmic transport of specific macromolecules while impeding the exchange of unsolicited material. However, key aspects of this gating mechanism remain controversial. To address this issue, we determined the nanoscopic behavior of the permeability barrier directly within yeast *S. cerevisiae* NPCs at transport-relevant timescales. We show that the large intrinsically disordered domains of phenylalanine-glycine repeat nucleoporins (FG Nups) exhibit highly dynamic fluctuations to create transient voids in the permeability barrier that continuously shape-shift and reseal, resembling a radial polymer brush. Together with cargo-carrying transport factors the FG domains form a feature called the central plug, which is also highly dynamic. Remarkably, NPC mutants with longer FG domains show interweaving meshwork-like behavior that attenuates nucleocytoplasmic transport *in vivo*. Importantly, the *bona fide* nanoscale NPC behaviors and morphologies are not recapitulated by *in vitro* FG domain hydrogels. NPCs also exclude self-assembling FG domain condensates *in vivo*, thereby indicating that the permeability barrier is not generated by a self-assembling phase condensate, but rather is largely a polymer brush, organized by the NPC scaffold, whose dynamic gating selectivity is strongly enhanced by the presence of transport factors.

## Introduction

Intrinsically disordered proteins (IDPs) confer dynamic qualities to higher order assemblies in cells whose structures remain challenging to resolve (Lyon et al., 2021). This problem is exemplified in nuclear pore complexes (NPCs), which are massive macromolecular assemblies that mediate the rapid bidirectional traffic of signal-specific cargoes entering or exiting the cell nucleus (Wing et al., 2022). Each NPC is assembled from multiple copies of thirty proteins termed nucleoporins (Nups) that form an eightfold rotationally symmetric core scaffold that can expand between 40 and 60 nm in diameter (Akey et al., 2022; Allegretti et al., 2020; Bley et al.; Fontana et al.; Kim et al., 2018; Mosalaganti et al.; Petrovic et al.; Schuller et al., 2021; Zhu et al.; Zimmerli et al., 2021). The core scaffold surrounds a central transport conduit that we term the central transporter (CT) (Akey, 1990; Akey *et al*., 2022; Kim *et al*., 2018). To facilitate nucleocytoplasmic transport, each NPC employs a subgroup of eleven Nups whose large intrinsically disordered, phenylalanine-glycine (FG)-repeat-rich domains (henceforth termed FG domains) emanate into the CT from their anchor sites at the NPC scaffold (Rout et al., 2000). Importantly, the FG domains generate a permeability barrier with two essential functions. First, cargo-carrying transport factors (TFs) engage in multivalent interactions with the FG repeats to selectively traverse the NPC (Bayliss et al., 2000; Hough et al., 2015; Kapinos et al., 2014; Milles et al., 2015). Second, the permeability barrier guards against non-specific molecules from gaining access into the cell nucleus (Popken et al., 2015; Timney et al., 2016). Disruptions to the NPC transport machinery are associated with diseases ranging from viral infections (Shen et al., 2021), neurodegeneration (Coyne and Rothstein, 2022) and cancer (Cagatay and Chook, 2018).

Despite recent advances in NPC structure refinement (Akey *et al*., 2022; Allegretti *et al*., 2020; Kim *et al*., 2018; Schuller *et al*., 2021; Zimmerli *et al*., 2021), the CT is generally excluded from cryo-electron microscopy (EM)-derived maps and the NPC permeability barrier remains unresolved. For the most part, it remains a challenge to visualize the disordered FG domains within the nanoscopic interior of the NPC. This is evident with respect to the presence of a denser amorphous feature called the central plug (CP) that is present in a subset of CTs and is thought to comprise of TFs and their cargoes in transit (Stoffler et al., 2003). Inevitably, NPC barrier models are informed by the *in vitro* “flavors” of purified FG domains that are suggested to consist of cohesive low charge content “GLFG” domains and repulsive high charge content “FxFG” domains (Yamada et al., 2010) (**Table S1**).

A great deal of work has demonstrated the ability for the FG domains to form *in vitro* assemblies, but these show a spectrum of behaviors that span different length scales depending on experimental design (Hoogenboom et al., 2021; Lemke, 2016; Schmidt and Gorlich, 2016; Zilman, 2018). These range from surface-tethered polymer brushes (Kapinos *et al*., 2014; Lim et al., 2007; Lim et al., 2006; Schoch et al., 2012; Wagner et al., 2015) to biomolecular condensates that can phase separate into liquid-like droplets (Celetti et al., 2020), amyloid fibrils (Ader et al., 2010; de Opakua et al., 2022; Halfmann et al., 2012) or hydrogels (Frey and Gorlich, 2007; 2009; Frey et al., 2018; Frey et al., 2006; Hülsmann et al., 2012; Labokha et al., 2013; Milles et al., 2013; Schmidt and Gorlich, 2015). To be precise, biomolecular condensates self-assemble by invoking cohesive inter-domain interactions to exclude nonspecific macromolecules. In comparison, polymer brushes exclude nonspecific macromolecules by entropic and excluded-volume (repulsive) interactions that can counteract attractive contributions between adjacent molecules because of spatial constraints imposed by surface tethering (Zhao and Brittain, 2000). Hence, two basic characteristics distinguish biomolecular condensates and polymer brushes: (i) the ability to self-assemble in the absence or presence of an anchoring scaffold, and (ii) the dynamic motion of the FG domains. An appreciation of these contextual differences has motivated varied efforts to examine the authentic behavior of the FG domains within the NPC (Atkinson et al., 2013; Cardarelli et al., 2012; Ma et al., 2016; Mattheyses et al., 2010; Mohamed et al., 2017; Sakiyama et al., 2016; Stanley et al., 2019; Stanley et al., 2018; Yu et al., 2022).

Several features make high-speed atomic force microscopy (HS-AFM) imaging advantageous for studying these dynamic biomolecular processes in aqueous physiological environments (Ando et al., 2013). First, the three-dimensional topographic resolution of HS-AFM is typically 2 to 3 nm in the XY plane and 0.15 nm in the Z direction. Second, HS-AFM uses a rapidly oscillating nanometer-sharp tip to tap on a surface intermittently at microsecond pulses (i.e., MHz frequencies) and sub-50 pN forces. This minimizes disturbances to a specimen as the impulse or energy being transferred is negligibly small (Ando, 2018). Third, molecular motion is captured at scan speeds of ∼100 milliseconds per image frame (Kodera et al., 2010), which falls within the range of NPC transport timescales that span between ∼10 ms for nuclear localization signal (NLS)-cargo and ∼200 ms for mRNA (Dange et al., 2008; Grünwald and Singer, 2010; Yang and Musser, 2006). In this way, HS-AFM has the power to resolve the dynamic motion of individual intrinsically disordered protein molecules (Kodera et al., 2021) and, as previously shown, is able to resolve the nanoscale dynamic behavior of FG domains in *X. laevis* oocyte NPCs (Sakiyama *et al*., 2016). In this study, we combined video-rate HS-AFM imaging with *in vivo* transport assays to address the nature of the permeability barrier *in situ* within native and mutant yeast NPCs.

## Results

### High-speed AFM analysis of isolated yeast NPCs

Isolated native yeast (WT) NPCs retain an intact structure and represents the contracted form of the normal range of NPC diameters (Akey *et al*., 2022; Allegretti *et al*., 2020; Kim *et al*., 2018; Zimmerli *et al*., 2021). This makes them excellent specimens in the current study, as their relatively uniform size minimizes potential variabilities in pore-to-pore behavior. Moreover, removing the nuclear envelope (NE) eliminates any membrane undulations that might influence dynamic FG domain behavior (Stanley *et al*., 2019). The CT diameter in freshly isolated WT NPCs was verified by low magnification HS-AFM scans to be 39.7 ± 6.1 nm consistent with the cytoplasmic face (Akey et al., 2022; Kim et al., 2018) where a majority bore 26.4 ± 8.0 nm-diameter CPs (termed +CP WT NPC) (**Fig. 1a and b; Fig. S1**). Further verification by negative stain EM showed that the overall size and eight-fold symmetric structure of +CP WT NPCs proved to be consistent with our published cryo-EM maps (Akey *et al*., 2022; Kim *et al*., 2018) (**Fig. S1**).

**Figure 1.**
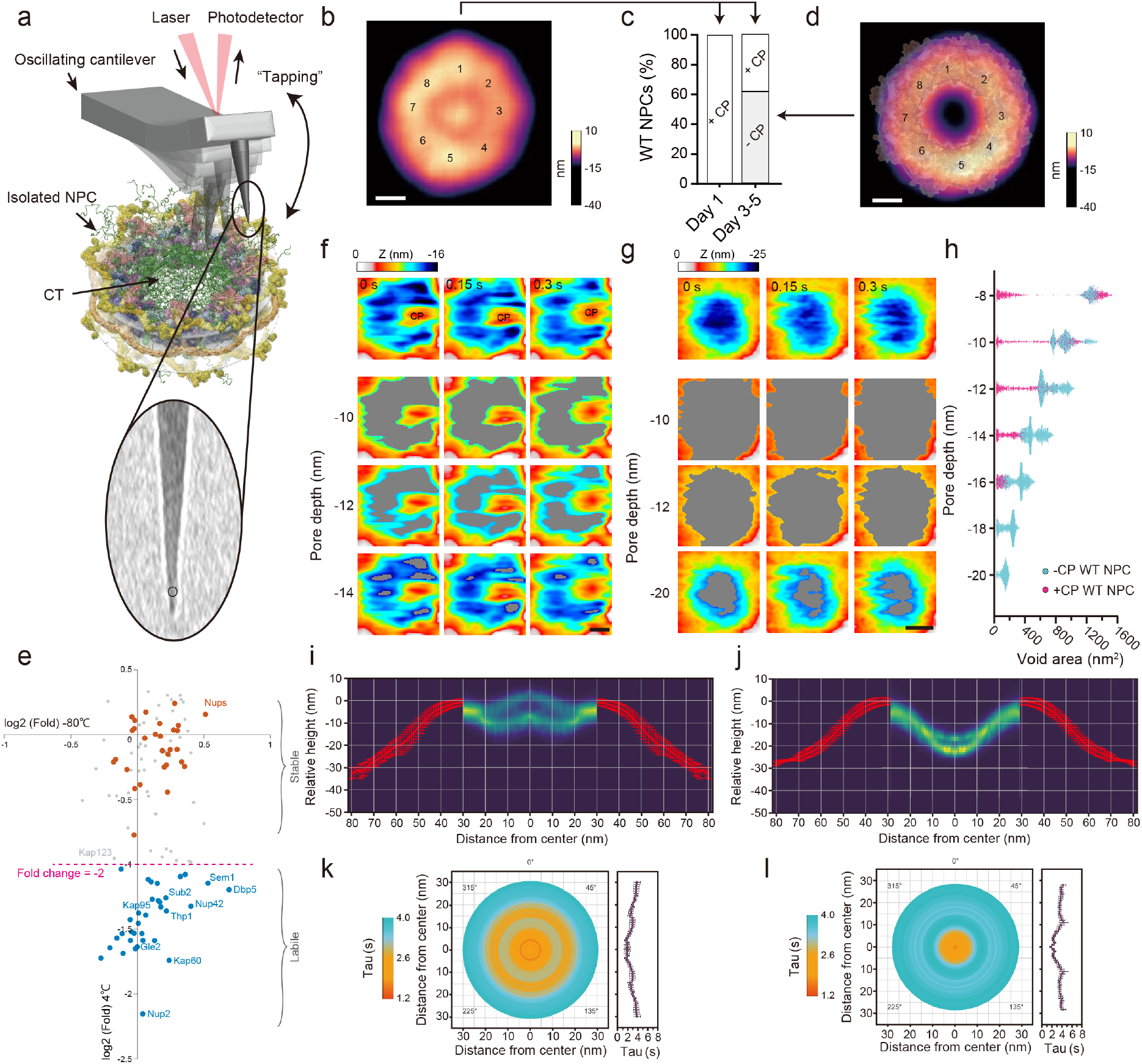
Direct visualisation of isolated yeast NPCs. **a**, Schematic description of the HS-AFM experiment to visualise the permeability barrier in isolated yeast NPCs. CT = central transporter. Inset: Transmission electron microscopy reveals the ∼2 nm-radius (circle) of a pristine HS-AFM tip. **b,** Average HS-AFM image resolves the CP and eightfold rotational symmetry (numbered) of +CP WT NPCs (n = 14). Scale bar, 20 nm. **c,** Percentage distribution of +CP and -CP WT NPCs (Day 1, n = 41; Day 3 to 5, n = 47). **d,** Integrative model structure of the isolated NPC (Kim et al., 2018) superimposed onto an average HS-AFM image of -CP WT NPCs (n = 15) emphasising their close agreement. Numbers indicate the eightfold rotational symmetry of the NPC scaffold. Scale bar, 20 nm. **e,** Comparison between the proteins associated with affinity-purified isolated yeast NPCs (Kim et al., 2018) incubated at 4°C for 5 days or snap-frozen in liquid nitrogen and stored at -80°C for 5 days as determined by label-free mass spectrometry quantification. Proteins showing a level of depletion higher than 2 fold in the 4°C stored NPCs are indicated in blue (proteasome components not labeled). Biological replicas n = 2. **f-g, Top row:** Consecutive HS-AFM images obtained at 0.15 s/frame resolves dynamic behavior in +CP WT NPCs **(f)** and -CP WT NPCs **(g)**. **2^nd^ to 4^th^ row**: Sequence of threshold images showing height-specific topographic features and voids (gray) that shape-shift over time at three different pore depths. Images in the 4th row represent the deepest accessible depths in the corresponding frames. CP = central plug; Scale bars, 10 nm. **h,** Scatter plot showing the relationship between pore depth and void area for +CP WT NPCs (n = 3, no. of frames ≈ 200 per NPC) and -CP WT NPCs (n = 3, no. of frames ≈ 600 per NPC). **i-j,** Peak-to-peak fluctuations in the Z-axis (turquoise) of +CP (n = 10) **(i)** and -CP WT NPCs (n = 19) **(j)** superimposed with their corresponding average NPC cross-sections (red). **k-l, Left:** Spatiotemporal dynamic map of the permeability barrier in +CP WT NPCs (n = 10, 3093 total frames analysed) **(k)** and -CP WT NPCs (n = 19, 7572 total frames analysed). **Right:** Plot of decay time (Tau) as a function of distance from the pore center. The light blue and orange colours correspond to Tau values that indicate ‘slower’ and ‘faster’ dynamics, respectively as shown in the corresponding lookup table. Data are mean ± s.e.m.

### Transport factors account for the main components of the central plug

NPCs are compositionally dominated by transport flux (Kim *et al*., 2018). *In situ* this includes a long-lived fraction of TFs such as Kapβ1 (human Kap95) (Kapinos et al., 2017; Lowe et al., 2015) whose slow rate of unbinding from the FG domains has been measured to be *K_off_* = 10^-5^ to 10^-6^ s^-1^ (Kapinos *et al*., 2014). But because permeability barrier models are generally premised on *in vitro* purified FG domains, we found it pertinent to examine *in situ* FG domain behavior in isolated NPCs depleted of CPs. We therefore took advantage of the selective exchange of TFs that occurs in isolated NPCs over time (Hakhverdyan et al., 2021) and found that their incubation in buffer was sufficient to facilitate TF-cargo depletion. After 72 hrs, 60% of isolated NPCs were depleted of any visible CPs (-CP WT NPC) being consistent with the rate of unbinding (Kapinos *et al*., 2014), to give an average CT diameter of 33.8 ± 4.1 nm that was unchanged from +CP WT NPCs (**Fig. 1c and d; Fig. S1**).

We assessed CP depletion by label-free quantitative mass spectrometry, which revealed that TFs and accessory proteins were indeed quantitatively depleted in comparison to freshly isolated NPCs (**Fig. 1e**). Indeed, a majority of the most abundant import factor, Kap95 and its adaptor Kap60 were displaced from -CP WT NPCs. Other TFs such as Kap123 also showed reduced levels. Thus, the CP is strongly associated with the presence of transiting TFs and cargoes in the NPC. In comparison, the abundant and dynamic mRNA export factors Mex67, Mtr2 or Yra1 were not significantly depleted (Hakhverdyan *et al*., 2021; Kim *et al*., 2018). These reside preferentially at the NPC periphery during transport (Ben-Yishay et al., 2019; Derrer et al., 2019), suggesting a relatively limited contribution to the CP. Reduction was also observed for proteasome components, mRNP remodeling factors and members of the Tho/TREX complex, along with Nup2 and Nup42, which are non-essential, peripheral FG Nups that are known to rapidly exchange *in vivo* (Hakhverdyan *et al*., 2021; Strawn et al., 2004). Importantly, the quantities of all other NPC components were not significantly depleted within the uncertainty of the method for the duration of our experiments. We further note that the samples for mass spectrometry naturally include partially assembled or disassembled NPCs, which as they would not appear as intact NPCs were excluded from HS-AFM analyses and could contribute to the loss of some materials.

### FG domains fluctuate radially towards the central plug

Dynamic molecular motion was captured using HS-AFM in both +CP WT NPCs and -CP WT NPCs by zooming into the CT of intact NPCs. To enable comparisons between data sets, we assigned a zero-height value to the highest feature of the NPC rim in zoomed out images as a point of reference and aligned subsequent zoomed in images to it (**Fig. S2**). Moreover, a constant imaging force of ∼40 pN was applied to all specimens and experimental conditions so that the resulting height data could be used as a proxy to gauge the range and strength of the permeability barrier.

In +CP WT NPCs, video playback revealed rapid fluctuations within the CT (**Fig. 1f; Movie S1**). Importantly, the overall morphologies and dynamic behaviors of the CT and CP in the isolated *S. cerevisiae* NPCs were entirely consistent with our previous observations in *X. laevis* oocyte NPCs (Sakiyama *et al*., 2016). This commonality prevails even though the organisms are divergent, and the isolation methodologies have little in common. As both preparations contain intact NPCs, and indeed (i) the oocyte preparation with its intact NE has been shown to be structurally intact (Eibauer et al., 2015) and fully transport-competent (Siebrasse and Peters, 2002) and (ii) the isolated yeast NPCs have been shown to be competent for rapid nuclear transport factor exchange (Hakhverdyan *et al*., 2021), these results strongly support the *in vivo* relevance of our observations.

IDPs show characteristic long-range conformational chain dynamics that span timescales of between 100 ns and 1 ms (Chowdhury et al., 2023). Effectively the dynamic “fuzzy” motion of the FG domains is downsampled in HS-AFM measurements, which we tested by simulating the HS-AFM experiment using a Brownian dynamics computational model of the NPC; this also confirmed that the HS-AFM sampling rate yielded information relevant to the pertinent time scales (**Fig. S3; Movies S2 and S3**). Interestingly, the FG domains display radial fluctuations in +CP WT NPCs that are drawn preferentially towards the CP, continuously colliding into and breaking contact with it. We suggest that this behavior sustains the dynamic movements of the CP, which physically manifests from the weak multivalent interactions between the FG repeats and TFs (Hayama et al., 2018; Hough *et al*., 2015; Kapinos *et al*., 2014; Milles *et al*., 2015; Raveh et al., 2016; Schoch *et al*., 2012; Wagner *et al*., 2015).

### Shape-shifting voids create dynamic passages to the NPC midplane

Strikingly, the radial fluctuations of the FG domains in +CP WT NPCs carve out transient voids that dynamically shape-shift and reseal within the permeability barrier. Moreover, these voids appear to be arranged in a roughly circular pattern around the CP. From the topographic information contained in HS-AFM images we quantified the size of the voids as a function of depth, being -16 nm at the deepest accessible point (**Fig. 1f** **and Fig. S4**). Here, the CP features prominently at the entrance of the CT whereas FG domain fluctuations are sparse. Only at a depth of ∼-10 nm do the FG domain fluctuations appear and are seen to extend across to the CP at -14 nm, being consistent with FG Nup bridges that link the scaffold to the CT (Akey *et al*., 2022; Kim *et al*., 2018). Consequently, the presence of these structures hindered the HS-AFM tip from probing deeper into the permeability barrier of +CP WT NPCs.

HS-AFM tips penetrated more deeply into -CP WT NPCs and gained access to depths of ∼-20 nm approaching the NPC midplane (Akey *et al*., 2022; Kim *et al*., 2018). Between -15 to -20 nm the -CP WT NPCs exhibit rapid FG Nup fluctuations that continuously extend and retract from the scaffold but did not coalesce nor span across the CT (**Fig. 1g** **and Fig. S4; Movie S4**). Above these depths, large voids fill the space and the CT is mostly unobstructed. The size of these voids decreases with depth but are generally larger than +CP WT NPCs (**Fig. 1h**). The average size of the lowest accessible voids at -20 nm was 134.5 ± 39.9 nm^2^, which can be approximated as circular holes bearing radii of between 5.5 nm and 7.5 nm. In contrast, the deepest accessible voids in +CP WT NPCs were shallower at -16 nm and were on average 85.4 ± 39.3 nm^2^, being equivalent to holes bearing radii of between 3.8 nm and 6.3 nm. NPCs exhibit the characteristics of a soft permeability barrier *in vivo*, where a continuously variable pore size allows for a continuous rather than discontinuous passive size exclusion relationship (Timney, 2016; Popken, 2016). Hence, this view is supported and rationalized by the appearance of shape-shifting voids, which show a large distribution of sizes particularly near the NPC midplane (**Fig. 1f-h**).

### The presence of the central plug accentuates dynamic behavior

Next, we adapted an autocorrelation function (ACF) analysis workflow for HS-AFM/AFM data (Kodera *et al*., 2021; Stanley *et al*., 2019) to generate spatiotemporal maps that characterize dynamic regions within the CT (**Fig. S5**). Briefly, this includes (i) constructing radial kymographs; (ii) extracting Z-fluctuations and computing ACFs from all kymographs; (iii) calculating the time-lag Tau, which is used as a readout for dynamic behavior; and (iv) determining average Tau values from multiple NPCs (**Fig. S6**). To determine the noise threshold in our experiments we further carried out control measurements on bare mica surfaces, which gave root-mean-square (RMS) values of 0.13 nm while holey defects in lipid bilayers and -CP WT NPCs gave RMS values of 0.35 nm and 2.8 nm, respectively (**Fig. S7**).

Superimposing all Z-fluctuations onto the average NPC height profile reveals the range of the permeability barrier which protrudes out of the CT in +CP WT NPCs but sags into the CT in -CP WT NPCs (**Fig. 1i and j**). Clearly, dynamic behavior (less correlated; smaller Tau) dominates in +CP WT NPCs and -CP WT NPCs with the former showing a larger dynamic zone than -CP WT NPCs (**Fig. 1k and l**). This large dynamic zone in +CP WT NPCs pertains to the movements of the CP and the FG domains that flank it; whereas, in -CP WT NPCs, the most dynamic zone derives from FG domain fluctuations that fall within a 10 nm-radius from the pore axis. Fanning further outwards towards the CT periphery sees decreased dynamics (more correlated; larger Tau) in both +CP WT NPCs and -CP WT NPCs due to the presence of the scaffold and the anchoring of the FG Nups to it, which naturally reduces their motion. Importantly, these results collectively agree with fluorescence-based measurements of dynamic FG Nup behaviour in intact NPCs *in vivo* (Atkinson *et al*., 2013; Yu *et al*., 2022) and in silico (Moussavi-Baygi and Mofrad, 2016; Winogradoff et al., 2022), and the restricted motion of FG domains near their anchor sites as inferred from cryo-EM data (Akey *et al*., 2022; Kim *et al*., 2018).

### GLFG Nups in maximal deletion-FG mutant NPCs do not form a crosslinked meshwork

It has been shown that it is possible to genomically delete the FG domains of several Nups in *S. cerevisiae*, and combine these deletions to eliminate more than 50% of the FG mass *in toto* within *in vivo* NPCs without loss of viability (Adams et al., 2016; Strawn *et al*., 2004). We used HS-AFM to examine two complementary mutants that have large changes in the FG composition of NPCs and are opposite in mass and FG flavor balance. First, we isolated septuple *nup42ΔFG nup159ΔFG nup60ΔFxF nup1ΔFxFG nup2ΔFxFG nsp1ΔFGΔFxFG* mutant NPCs (Strawn *et al*., 2004) (SWY3062; abbreviated ΔFG NPC), which reduced the theoretical FG repeat concentration in the NPC to ∼6 mg/ml (∼27 mM); 51% below the estimated FG repeat concentration in WT NPCs of ∼53 mM (Akey *et al*., 2022; Kim *et al*., 2018) (**Table S2**). Importantly, ΔFG NPCs represent a maximal FG-deletion mutant where all FxFG domains are purged but nevertheless retain the size, inner diameter, overall morphology and non-FG containing Nup composition of WT NPCs (**Fig. S1 and Fig. S8**). Moreover, in both this and a second mutant (below) the FG domain fluctuations were significantly different in comparison to WT NPCs, further underscoring that the fluctuations observed by HS-AFM are indeed the FG domains. As many of the remaining FG domains, namely Nup100, Nup116, Nup49, Nup57

and Nup145N contain the most “cohesive” GLFG domains, we asked if this composition of FG domains might form a pore-spanning meshwork within the CT as predicted by the selective phase model (Frey and Gorlich, 2007; 2009; Frey *et al*., 2006; Hülsmann *et al*., 2012; Labokha *et al*., 2013; Schmidt and Gorlich, 2015).

Freshly isolated ΔFG NPCs featured CPs (+CP ΔFG NPCs) that were significantly less mobile in comparison to WT NPCs. Moreover, the CP is surrounded by FG domains that fluctuate in place to expose voids that undergo minimal shape changes (**Fig. 2a** **and Fig. S4; Movie S5**) thereby suggesting that the mobility of the CP in WT NPCs may be enhanced by dynamic FxFG domains (Otto et al., 2023). When examining -CP ΔFG NPCs, the GLFG domains do not form a pore-spanning meshwork, but rather exhibit smaller fluctuations that surround the inner scaffold surface. Collectively, this resembles an annulus-like layer that constricts the CT (**Fig. 2b** **and Fig. S4; Movie S6**). For this reason, the overall void size in -CP ΔFG NPCs was consistently smaller than -CP WT NPCs (**Fig. 2c**) even though 51% of the total FG mass was missing from -CP ΔFG NPCs. Descending towards the midplane, the CT narrows into a funnel-like constriction that exposes a singular void at the deepest accessible depth of -18 nm. Its average size is 61.5 ± 21.2 nm^2^, which is approximately a circular hole of 3.6 to 5.1 nm in radius. Indeed, a high-density ring-like barrier is predicted to form at the midplane when FxFG domains are absent (Ghavami et al., 2014; Huang et al., 2020). Overall, these behaviors account for the slower dynamics and reduced Z-fluctuations in ΔFG NPCs (**Fig. 2d to g**).

**Figure 2.**
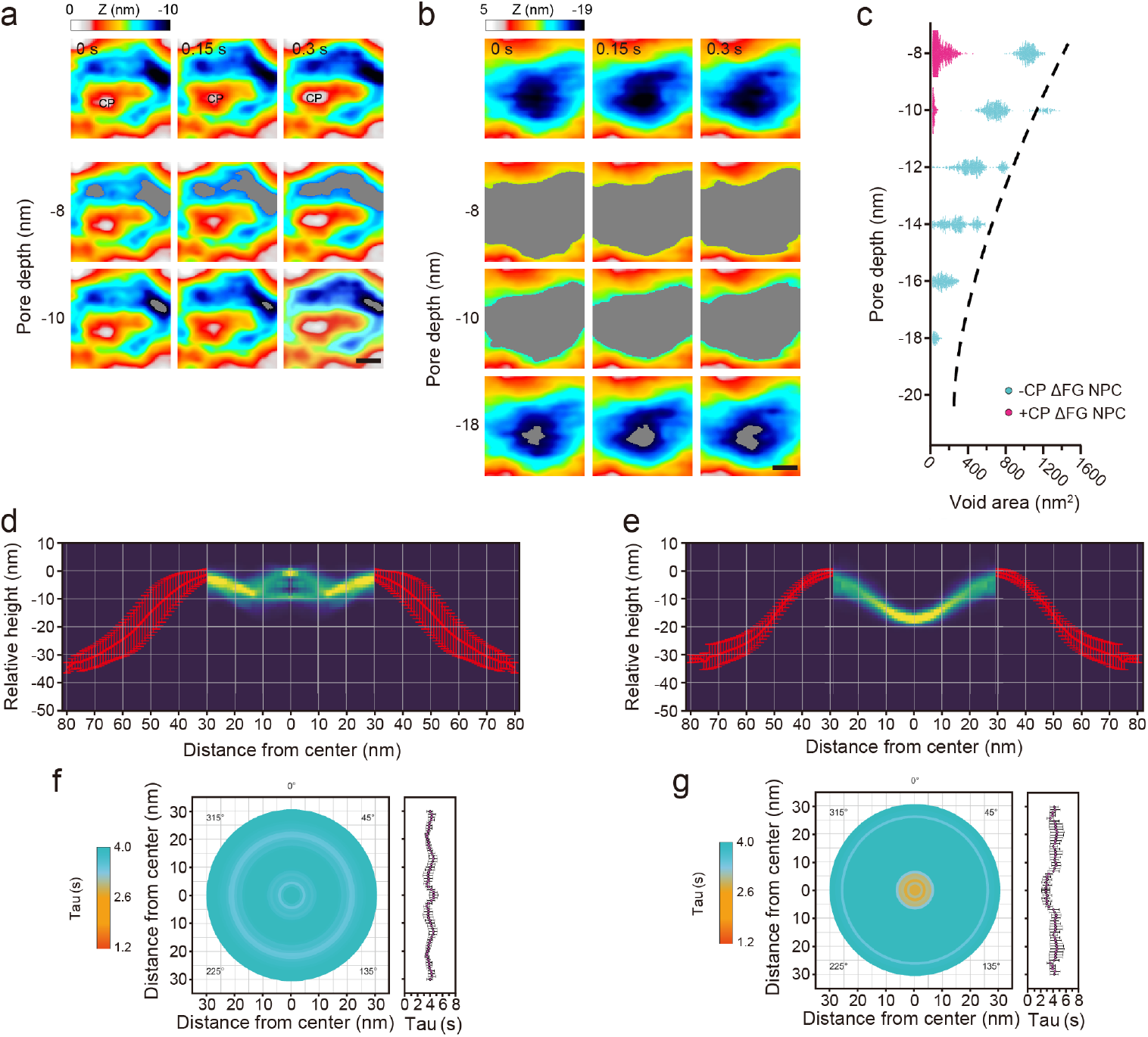
Dynamics of the permeability barrier in ΔFG NPCs. **a-b**, **Top row:** Consecutive HS-AFM images obtained at 0.15 s/frame resolves dynamic behavior in +CP ΔFG NPCs **(a)** and -CP ΔFG NPCs **(b)**. **2^nd^ to 3^rd^/4^th^ row**: Sequence of threshold images showing height-specific topographic features and voids (gray) that shape-shift over time at two **(a)** or three **(b)** different pore depths. Images in the 3^rd^ or 4^th^ row represent the deepest accessible depths in the corresponding frames. CP = central plug; Scale bars, 10 nm. **c,** Scatter plot showing the relationship between pore depth and void area for +CP ΔFG NPCs (n = 3, no. of frames ≈ 600 per NPC) and -CP ΔFG NPCs (n = 3, no. of frames ≈ 400 per NPC). The dashed line denotes the largest void areas in -CP WT NPCs and is included as a guide for comparison. **d-e,** Peak-to-peak fluctuations in the Z-axis (turquoise) of +CP ΔFG NPCs (n = 8) **(d)** and - CP ΔFG NPCs (n = 8) **(e)** superimposed with their corresponding average NPC cross-sections (red). **f-g, Left:** Spatiotemporal map of the permeability barrier in +CP ΔFG NPCs (n = 8, 4426 total frames analyzed) **(f)** and -CP ΔFG NPCs (n = 8, 3300 total frames analyzed) **(g)**. **Right:** Plot of decay time (Tau) as a function of the distance from the pore center. The light blue and orange colours correspond to Tau values that indicate ‘slower’ and ‘faster’ dynamics, respectively as shown in the corresponding lookup table. Data are mean ± s.e.m.

Generally, polymer brushes form when the spacing between surface-tethered polymer chains is smaller than their hydrodynamic diameters (Zhao and Brittain, 2000). To verify this, we computed the average next-neighbor distance between anchor sites that project the FG domains into the CT based on our integrative model of isolated NPCs (Kim *et al*., 2018). This gave 5.3 nm for the cytoplasmic lobe, 3.5 nm at the central waist and 7.3 nm in the nucleoplasmic lobe (**Fig. S9**). Apart from Nup145, these values are smaller than the predicted hydrodynamic diameters of the FG repeat domains found in each location (Yamada *et al*., 2010) (**Table S1**) and satisfy the requirement for polymer brush formation in the WT NPC. Then by omitting Nup42, Nup159, Nup60, Nup1, Nup2 and Nsp1, these values increased to 10.4 nm for the cytoplasmic lobe, 3.7 nm at the central waist and 14.5 nm for the nucleoplasmic lobe in ΔFG NPCs (**Fig. S9**). Clearly, the FG Nups in both cytoplasmic and nucleoplasmic lobes can no longer satisfy the requirement for full polymer brush formation, much less form a meshwork, being consistent with our current observations. Still, the grafting distance at the central waist in ΔFG NPCs is essentially unaltered with respect to WT NPCs, which might explain why their selective gating function is preserved (Strawn *et al*., 2004).

### Meshwork formation in double-length Nsp1 mutant NPCs

The previous results raised the issue that the FG repeat density in the NPC is balanced between being high enough to form a brush, but not so high as to engender other deleterious states. Therefore, we significantly *increased* the overall FG repeat amount and FG domain length in the most abundant FG Nup by generating a novel yeast strain containing double FG repeat domain-length Nsp1 constructs (aa1-591-2-565-592-end; termed Nsp1FGx2) (**Fig. S8**). This raised the theoretical FG repeat concentration in the NPC by 30% to ∼16 mg/ml (∼69 mM) over the estimated FG repeat concentration in WT NPCs (Akey *et al*., 2022; Kim *et al*., 2018) (**Table S2**). Remarkably, Nsp1FGx2 NPCs exhibited distinct hallmarks of meshwork formation. Chief among these was the presence of highly persistent hyper-elongated FG domain fluctuations, which coalesced with the CP in +CP Nsp1FGx2 NPCs (**Fig. 3a** **and Fig. S4; Movie S7**); or resembled interweaving meshworks that occluded the CT in -CP Nsp1FGx2 NPCs (**Fig. 3b** **and Fig. S4; Movie S8**). Voids also appeared at shallower depths of -6 nm and -12 nm in +CP Nsp1FGx2 NPCs and -CP Nsp1FGx2 NPCs, respectively (**Fig. 3c; Fig. S1**), despite bearing the same inner diameter as WT NPCs. Consequently, this resulted in a flatter distribution of Z-fluctuations in +CP Nsp1FGx2 NPCs and -CP Nsp1FGx2 NPCs, which were considerably less dynamic than WT NPCs (**Fig. 3d to g**). Taken together, our results indicate that the FG content in NPCs is tuned to a range that optimizes the formation of a polymer brush without entangling the FG domains into a meshwork.

**Figure 3.**
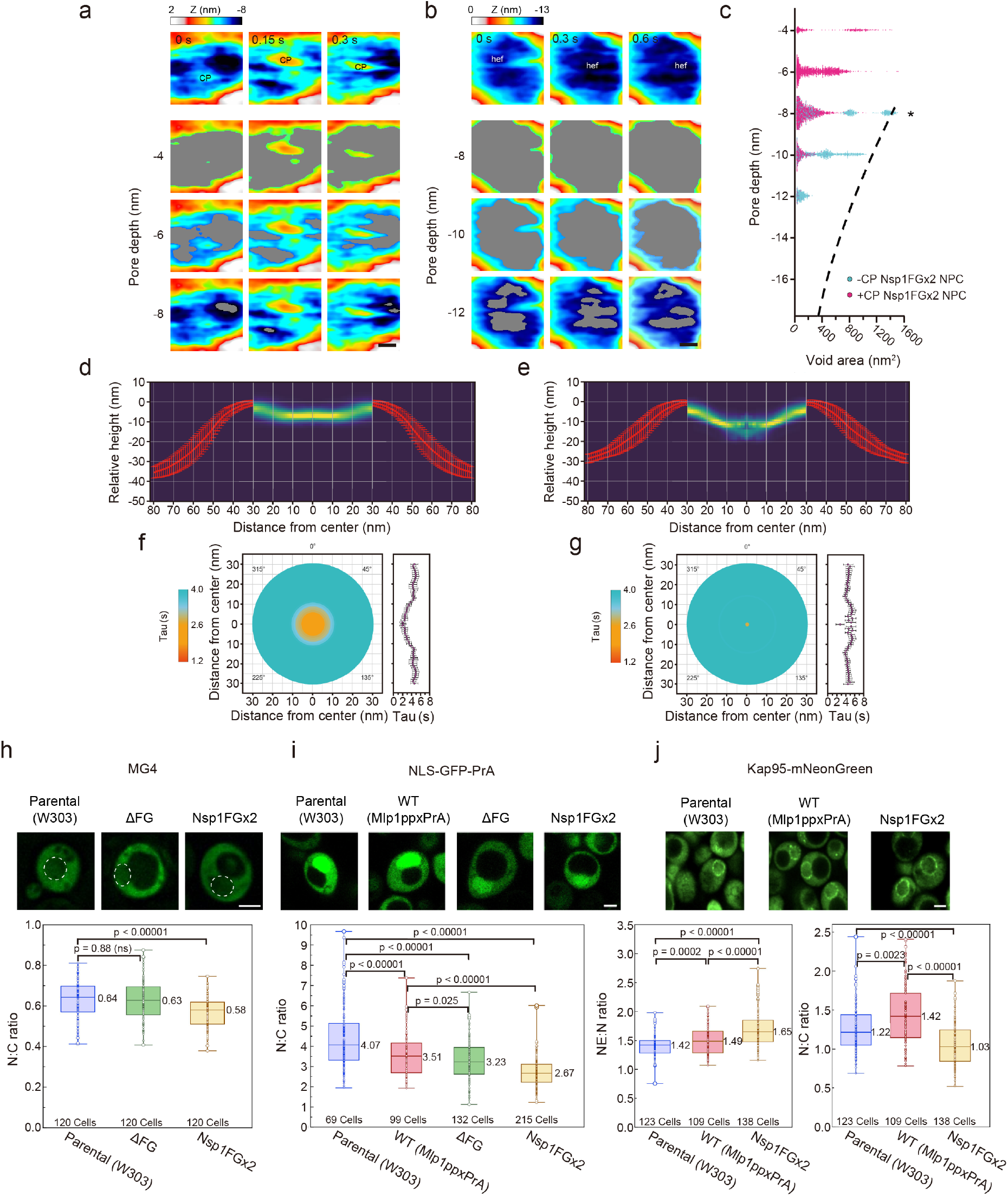
Nsp1FGx2 NPCs attenuate selective transport *in vivo.* **a-b**, **Top row:** Consecutive HS-AFM images obtained at 0.15 s/frame resolves dynamic behavior in +CP Nsp1FGx2 NPCs **(a)** and -CP Nsp1FGx2 NPCs **(b)**. **2^nd^ to 4^th^ row**: Sequence of threshold images showing height-specific features and voids (gray) that shape-shift over time at three different pore depths. Images in the 4^th^ row represent the deepest accessible depths in the corresponding frames. CP = central plug; hef = hyper-elongated fluctuations. Scale bars, 10 nm. **c,** Scatter plot showing the relationship between pore depth and void area for +CP Nsp1FGx2 NPCs (n = 4, no. of frames ≈ 447 per NPC) and -CP Nsp1FGx2 NPCs (n = 3, no. of frames ≈ 500 per NPC). The dashed line denotes the largest void areas in -CP WT NPCs and is included for comparison. Note: The highest features in -CP Nsp1FGx2 NPCs are resolved at a pore depth of -8 nm (⭑). **d-e,** Peak-to-peak fluctuations in the Z-axis (turquoise) of +CP Nsp1FGx2 NPCs (n = 6) **(d)** and -CP Nsp1FGx2 NPCs (n = 7) **(e)** superimposed with their corresponding average NPC cross-sections (red). **f-g, Left:** Spatiotemporal map of the permeability barrier in +CP Nsp1FGx2 NPCs (n = 6, 2912 total frames analyzed) **(f)** and -CP Nsp1FGx2 NPCs (n = 7, 3239 total frames analyzed) **(g)**. **Right:** Plot of decay time (Tau) as a function of the distance from the pore center. The light blue and orange colours correspond to Tau values that indicate ‘slower’ and ‘faster’ dynamics, respectively as shown in the corresponding lookup table. Data are mean ± s.e.m. **h-i,** Steady state nuclear localization of MG4 (**h**; no. of replicates = 4) and SV40NLS-GFP-PRA (**i**; no. of replicates = 4) quantified by its ratio of fluorescence intensity in the nucleus and cytoplasm (N:C ratio) in the indicated strains; NE-ER marker mCherry-L-TM is used to locate the nucleus in **(h)** (indicated by white dotted circles). **j,** N:C ratio of Kap95-mNeonGreen fluorescence intensity in the nucleus and cytoplasm (right) shown together with the ratio of its fluorescence intensity at the nuclear envelope and nucleus (left; NE:N ratio) in the indicated strains (no. of replicates = 4). Number of cells analyzed, median values, first and third quartiles are indicated in the box plots. Scale bars, 2 μm.

### Double-length Nsp1 mutants tighten passive permeability but impair selective transport *in* vivo

We next tested whether these mutations were indeed generating deleterious states in the NPCs. We asked how the observed differences in the selective permeability barrier of Nsp1FGx2, ΔFG NPCs and WT NPCs might affect NPC function and cell fitness. In comparison to the WT, phenotypic analyses showed a fitness defect in both mutants, milder in Nsp1FGx2 and more severe in ΔFG (**Fig. S8**). To test for differences in passive permeability, we expressed two non-specific reporters of different sizes: 2xgreen fluorescent protein (2xGFP) (54 kDa) (**Fig. S10**) and a maltose binding protein-4xGFP fusion protein (MG4; 150 kDa) (Popken *et al*., 2015) (**Fig. 3h**). Although the 2xGFP returned similar N:C ratios in all three strains due to its small size, we found a significant reduction in the N:C ratio of MG4 in Nsp1FGx2 cells compared to WT. Thereafter, we expressed a simian virus 40 nuclear localization signal (SV40NLS)-GFP-protein A (PrA) fusion reporter to test for changes in active transport along the Kap60-Kap95 pathway. Notably, nuclear accumulation of SV40NLS-GFP-PrA in Nsp1FGx2 cells was 24% less than in WT cells (**Fig. 3i**), thereby signifying a clear import defect. Meanwhile, ΔFG NPCs showed the same passive and selective permeabilities as WT NPCs being consistent with our previous findings (Popken *et al*., 2015; Timney *et al*., 2016) (**Fig. 3h and i**). Based on our analysis of grafting distance, the central waist in ΔFG NPCs is essentially unaltered with respect to WT NPCs (**Fig. S9**) so that this sufficiently maintains a permeability barrier comprised solely of GLFG domain brushes (**Fig. 2b and c**).

TFs enrich at NPCs and play a role in reinforcing the permeability barrier (Kalita et al., 2022; Kapinos *et al*., 2017; Kim *et al*., 2018). We therefore expressed Kap95-mNeonGreen in mutant strains and WT cells to ascertain its enrichment at the NPCs (**Fig. 3j**). The NE:N ratio of Kap95-mNeonGreen showed an 11% increase in enrichment in Nsp1FGx2 NPCs over WT, which is most likely caused by the increased concentration of FG repeats. However, this was accompanied by a 27% decrease in its N:C ratio, thereby suggesting that an oversaturation of Kap95-mNeonGreen at Nsp1FGx2 NPCs resulted in a decrease in translocation probability due to crowding (Zheng and Zilman, 2023). Indeed, this is commensurate with the reduction of SV40NLS-GFP-PrA in the nucleus (**Fig. 3i**). When combined, our results indicate that an overtightening of the passive permeability barrier - as a result of FG domain meshwork formation and an over-saturation of Kap95-mNeonGreen in Nsp1FGx2 NPCs – obstructs NLS-cargo delivery into the cell nucleus. This shows that FG domain meshwork formation significantly impairs nucleocytoplasmic transport.

### *In vitro* FG Nup gels are morphologically heterogeneous and display FG-gated “gel-holes”

Following from above, we wondered how the *in situ* characteristics of the NPC permeability barrier compared with those of *in vitro* FG condensates. Following established protocols (Schmidt and Gorlich, 2015), fluorescein-5-maleimide-labelled yNup100FG (the yeast homolog of human Nup98) phase separated into ∼10 µm sized particles *in vitro* that deviated slightly from a perfectly spherical shape as evidence of their solid hydrogel-like behavior, as described previously. As before, these yNup100FG hydrogel particles were selectively permeable to Kap95 but excluded Alexa647-labelled BSA passive reporters (**Fig. 4a**) (Frey and Gorlich, 2007; 2009; Frey *et al*., 2018; Hülsmann *et al*., 2012; Milles *et al*., 2013; Schmidt and Gorlich, 2015). Further photobleaching quantification of yNup100FG showed that the mobile fraction of each particle was ∼20% both in the absence and presence of Kap95 with the remaining fractions being immobile, which is consistent with previous qualitative observations (**Fig. 4b**) (Schmidt and Gorlich, 2015). Meanwhile, Kap95 partitioned selectively into the yNup100FG particles with ∼80% being in the mobile fraction.

**Figure 4.**
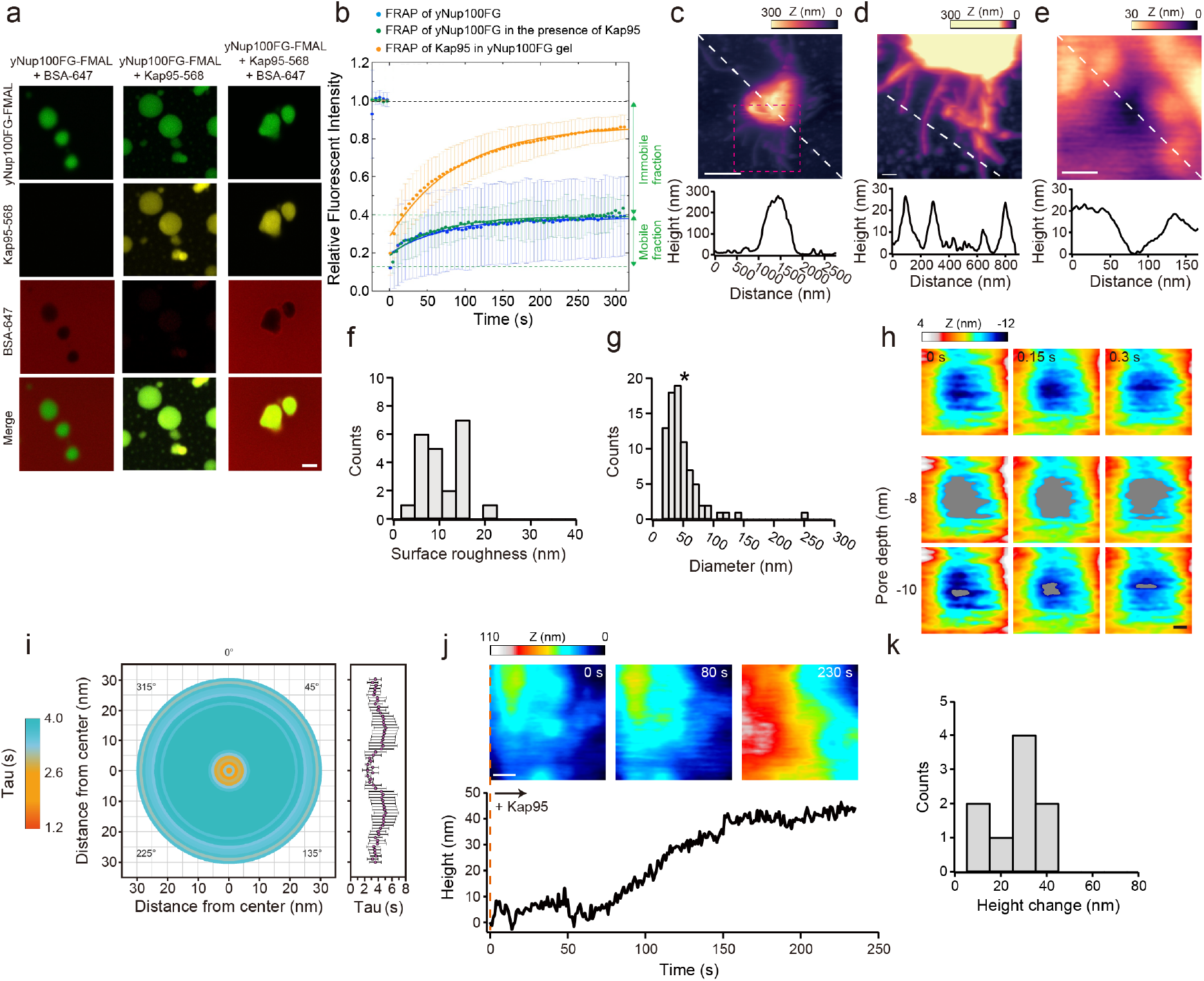
Nanoscale characterization of yeast Nup100FG hydrogels. **a,** yNup100FG labelled with fluorescein-5-maleimide (yNup100FG-FMAL) gels exclude passive BSA-AF647 molecules in the absence and presence of Kap95-AF568. Scale bar, 2 μm. **b,** Solid lines indicating best fits to the mean fluorescence recovery curves for yNup100FG-FMAL alone (cyan), yNup100FG-AF488 in the presence of Kap95-AF568 (green) and Kap95-AF568 in a yNup100FG gel (orange). Mobile fractions of yNup100FG in the absence and presence of Kap95 are 23 ± 10 % and 22 ± 4 % while their half-time recoveries are 69 ± 23 s (n = 11) and 60 ± 28 s (n = 3), respectively. In comparison, 81 ± 4 % of Kap95 in a yNup100FG gel is mobile with a half-time recovery of 65 ± 15 s (n = 4). Mobile fraction of yNup100FG is calculated from the first intensity value after photobleaching and the recovery plateau region. **c,** HS-AFM image and cross-section of a yNup100FG particle reveals its heterogeneous morphology. The cross-sectional profile corresponds to the white dashed line in the image. Scale bar, 500 nm. **d,** Zoom-in of the amyloid fibrils bounded by the magenta box in c. The cross-sectional profile corresponds to the white dashed line in the image and shows the thickness of each amyloid fibril. Scale bar, 100 nm. **e,** Closer inspection reveals the presence of granular aggregates that surround a gel-hole. The cross-sectional profile corresponds to the white dashed line in the image. Scale bar, 30 nm. **f**, Distribution of hydrogel surface roughness (n = 22; mean ± s.d. = 11.8 ± 4.5 nm). **g**, Distribution of gel-hole diameters (n = 82; mean ± s.d. = 52.4 ± 31 nm). Black star represents NPC diameter. **h,** Consecutive HS-AFM images obtained at 0.15 s/frame resolves dynamic fluctuations in a gel-hole. **2^nd^ to 3^rd^ row**: Sequence of threshold images showing height-specific topographic features and voids (gray) that shape-shift over time at two different pore depths. Images in the 3rd row represent the deepest accessible depths in the corresponding frames. Scale bar, 10 nm. **i,** Left: Dynamic map of the gel-holes (n = 7, 3727 total frames analyzed). Right: Plot of decay time (Tau) as a function of the distance from the pore center. **j,** Upper: Time-lapse HS-AFM images recorded at 1 s/frame showing an increase in height of the hydrogel during Kap95 binding. Lower: Corresponding plot of height as a function of time following the addition of 1 μM Kap95 at 0 s. **k,** Histogram of height changes at the hydrogel due to Kap95 binding. The average height increase is 31.2 ± 10.8 nm (n = 9).

Although the yNup100FG particles exhibited qualities that were consistent with previously reported behaviors that replicate certain selective permeability properties of the NPC (Frey and Gorlich, 2007; 2009; Frey *et al*., 2018; Hülsmann *et al*., 2012; Milles *et al*., 2013; Schmidt and Gorlich, 2015), we were puzzled as to how Kap95 could traverse them like a liquid even when the majority of yNup100FG was immobile. To explore this further, we used HS-AFM to resolve the morphology of the yNup100FG particles, at a spatial and temporal resolution commensurate with that of NPC behaviors (above) but one at which these hydrogels have not been analyzed previously. The hydrogel particles proved extremely morphologically heterogeneous at the nanoscale, contrasting with the high degree of organization seen within the CT of *in situ* NPCs. Granular aggregates and nanometer-sized crevices we term “gel-holes” dominated the particle surface, while ∼25 nm-thick amyloid-like fibrils protruded from the main body of the particle reaching lengths of over 1 µm (Halfmann *et al*., 2012; Milles *et al*., 2013; Schmidt and Gorlich, 2015) (**Fig. 4c to e**). Altogether these resulted in large variations in surface roughness that underscores the heterogeneity of the hydrogels (**Fig. 4f**). Surprisingly, the gel-holes spanned between 10 nm and 100 nm in diameter with many being coincident with NPC size (**Fig. 4g**), consistent with previous reports (Milles *et al*., 2013). Moreover, the gel-holes also contained FG repeat domain fluctuations and voids, which were significantly more dynamic than the main body of the hydrogel itself (**Fig. 4h and i**). At a depth of -10 nm, the average void size was 154.5 ± 66.3 nm^2^, which can be approximated as circular holes bearing 5.3 to 8.4 nm in radii. Hence, it is fortuitous that hydrogel-based estimates are congruent with the lower limit of passive diffusion in the NPC (Frey and Gorlich, 2007; 2009; Hülsmann *et al*., 2012; Labokha *et al*., 2013; Mohr et al., 2009; Schmidt and Gorlich, 2015), given that gel-holes share similar void sizes with WT NPCs.

Subsequently, adding Kap95 did not increase the number of nanosized holes as one would expect if individual crosslinks were being locally “unfastened” or “melted” as previously suggested (Frey and Gorlich, 2007; 2009; Hülsmann *et al*., 2012; Labokha *et al*., 2013; Schmidt and Gorlich, 2015). Instead, the height of the particle increased (**Fig. 4j and k**) due to the binding of Kap95 to surface-exposed yNup100FG repeats (Milles *et al*., 2013); similar to when Kaps bind to planar FG domain layers (Kapinos *et al*., 2014; Schoch *et al*., 2012; Vovk et al., 2016; Wagner *et al*., 2015). It thus appears that the gel-holes likely act as selective passageways for Kap95 into the particle interior and may account for much of the transport-like properties of the gel – although importantly, the condensate properties of the bulk of the gel do not appear to mimic NPC behavior at the nanoscale.

### NPCs resist fusion with Nup100FG liquid condensates *in vivo*

More recently, the model of a highly crosslinked FG hydrogel has been somewhat superseded by their suggestion of also forming liquid-liquid phase separation (LLPS)-like condensates, both *in vitro* (Celetti *et al*., 2020) and *in vivo* (Kuiper et al., 2022; Prophet et al., 2022). A basic and defining attribute of liquid-like biomolecular condensates lies in their ability to self-assemble and coalesce *in vivo* in the absence of an organizing scaffold (Brangwynne et al., 2009). Thus, NupFG liquid condensates should form at or fuse with NPCs if it presented a liquid-like permeability barrier *in vivo*. To test for such behavior, GFP-FG domains (Nup100FG, Nup116FG and hNup153FG), which have been described as cohesive and hydrogel / LLPS forming, and other GFP-FG domains not generally considered cohesive (Nup159FG, Nsp1FG and Nup60FG) were over-expressed for 1h in WT strains (**Fig. 5a** **and Fig. S11a**). Nup100FG, Nup116FG and hNup153FG indeed formed foci in the cytoplasm and nucleoplasm (**Fig. 5a**). Nup159FG, Nsp1FG and Nup60FG were found to localize throughout the cell without a defined pattern (**Fig. 5a**). Remarkably however, none of the GFP-FG domains showed a punctate nuclear rim staining pattern characteristic of NPC co-localization **(Fig. S11b)**, indicating they do not co-condense with the NPC. 90% of Nup100FG foci are solubilized by addition of 1,6 hexanediol indicating they are liquid-like condensates (Alberti et al., 2019; Elbaum-Garfinkle, 2019) (**Fig. S11c and d**). Individual Nup100FG condensates are localized at the cytoplasm, nucleoplasm or occasionally adjacent to the NE, which shows that the soluble GFP-FG constructs or condensates can both access and traverse the permeability barrier of WT and mutant NPCs without coalescing with it (**Fig. 5b**). This contrasted with controls where GFP-tagged full length FG Nups were expressed under their endogenous promoter and correctly localized to NPCs (**Fig. S11b**); thus, in the absence of a structured scaffold anchor, FG Nups will not concentrate or condense at NPC. This in turn indicates that the FG domains in the permeability barrier itself cannot nucleate a self-organized condensate.

**Figure 5.**
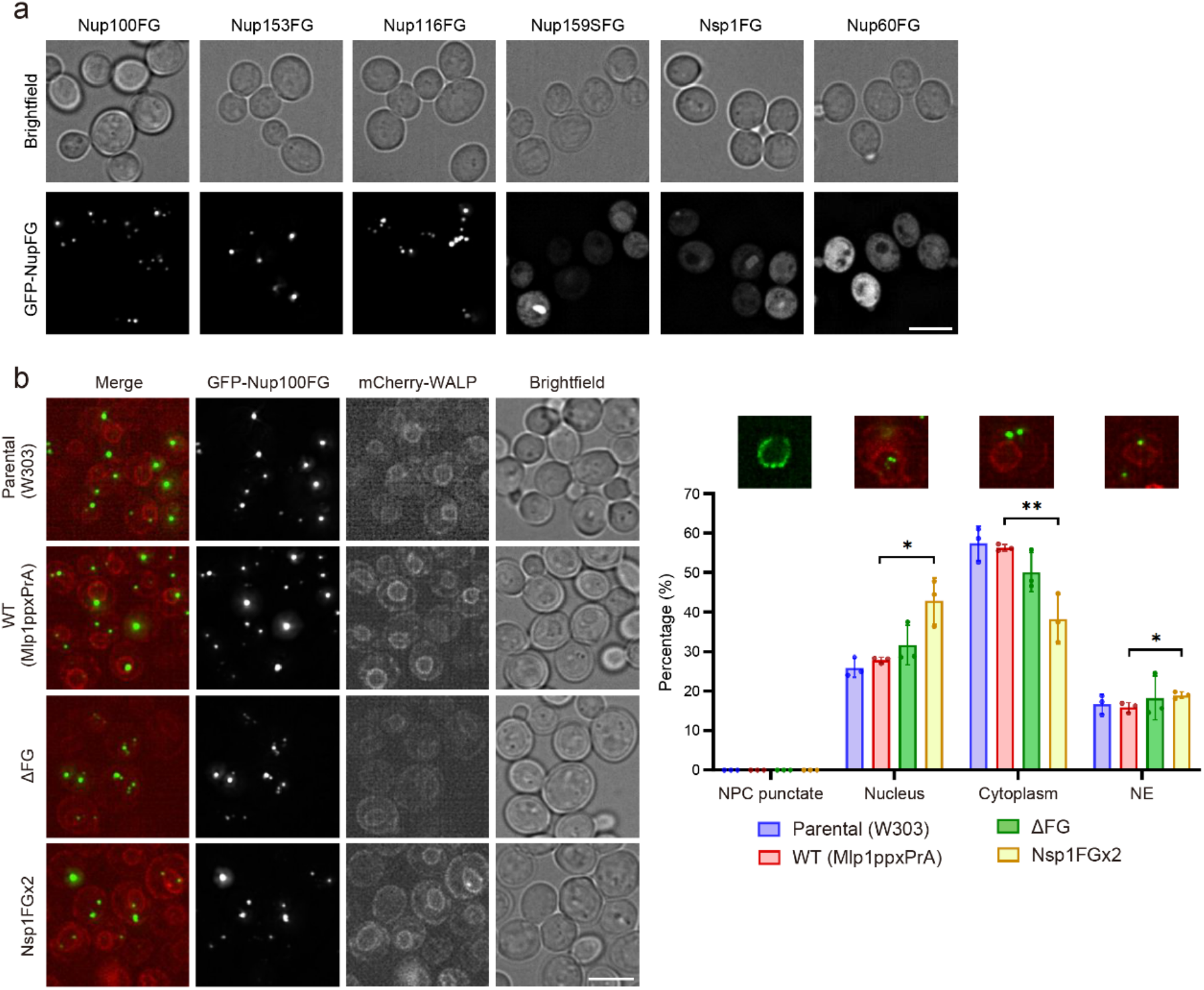
NPCs resist fusion with Nup100FG liquid condensates. **a**, Cellular localization of different FG domains tagged with GFP. Maximal projection of 4 z-stacks, 0,2 μm interval. Scale bar, 5 μm. See **Fig. S11a and b** for brighter images and full-length Nup controls, respectively. **b**, Cellular localization of the Nup100FG domain tagged with GFP, over-expressed for 1h in the indicated strains. Constitutively expressed mCherry-WALP-HDEL (red) is used as a NE-ER marker. Maximal projection of 4 z-stacks, 0,2 μm interval. Scale bar, 5 μm. Graph represents the quantification of GFP-Nup100FG localization (no. of replicates = 3; mean ± s.d.). Number of condensates analyzed in each replicate: 100-200. Exemplary images are shown; ‘NPC punctate’ depicts full length Nup100-GFP. See **Fig. S11c and d** for 1,6 hexanediol controls and expression levels, respectively.

### FG domain dynamics is conserved across species

Finally, we ranked the dynamics of the permeability barrier by plotting the average Tau values obtained from WT and mutant yeast NPCs along with the FG-gated gel-holes and their surrounding gel surfaces from yNup100FG particles (**Fig. 6a**). Additionally, we computed Tau from our previous *X. laevis* oocyte NPC data (Sakiyama *et al*., 2016). All NPCs were in comparable ranges, with the oocyte NPCs being the most dynamic followed by yeast WT, ΔFG and Nsp1FGx2 NPCs. This is consistent with the fact that Nup98 represents the only “cohesive” (or in our observations, least dynamic) GLFG Nup in vertebrate NPCs that are otherwise dominated by FxFG Nups. By comparison, the *in vitro* hydrogel particles were very different from the NPCs; though the FG-gated gel-holes exhibited dynamics that were intermediate between ΔFG NPCs and Nsp1FGx2 NPCs. The yNup100FG particle surfaces were the least dynamic, and as there must be free FG repeats at this surface this likely overestimates the actual mobility of the bulk condensate (e.g., as measured by fluorescence recovery, **Fig. 4b**). Hence, we conclude that the selective permeability of the yNup100FG particles likely derives from FG-gated gel-holes, and not its surrounding surfaces or gel-like substance.

**Figure 6.**
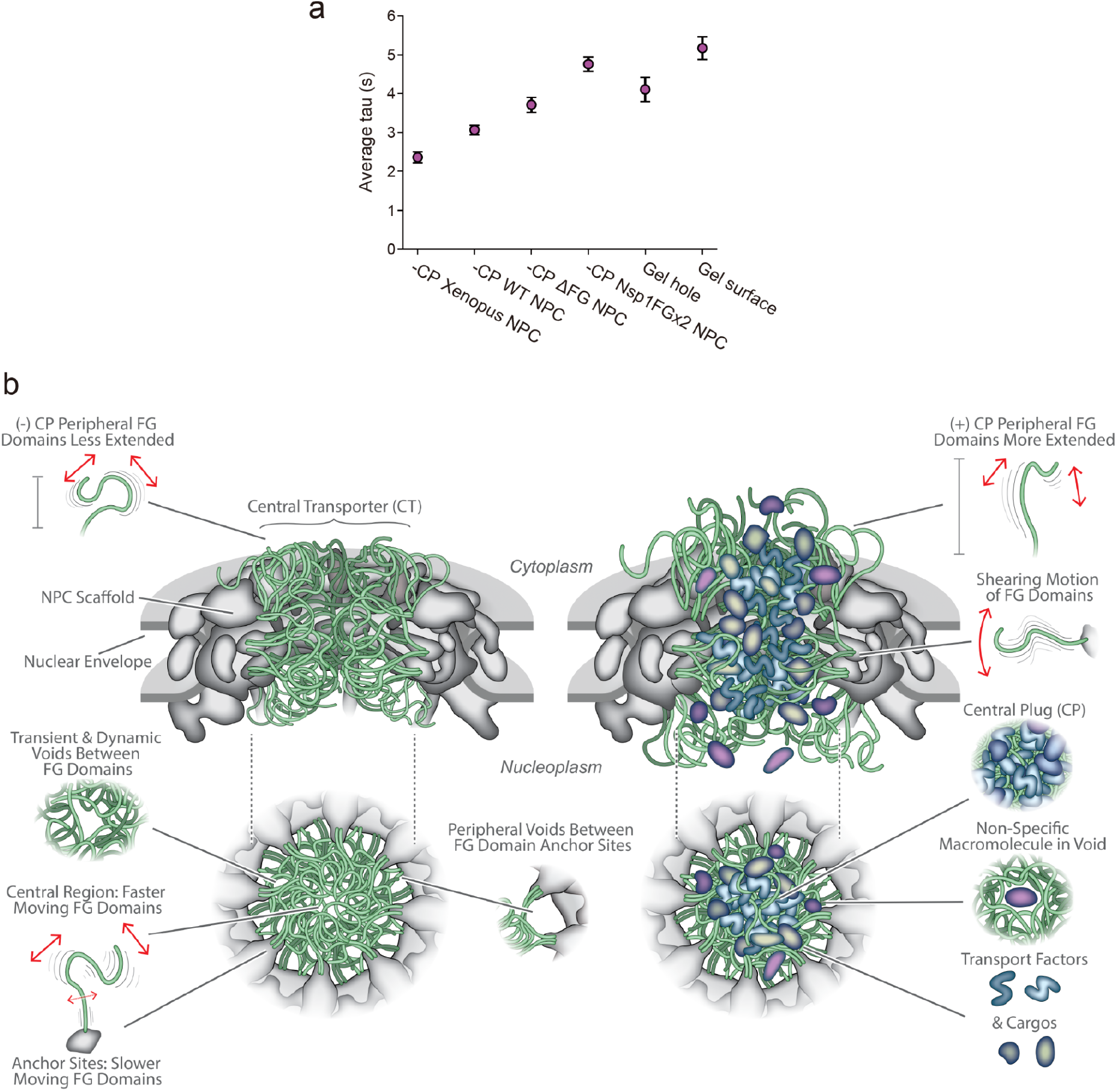
Dynamic molecular mechanism of the NPC permeability barrier. **a**, Plot of ascending Tau ranks the dynamics (fastest to slowest) of the permeability barrier across species and yeast strains and compares it with the FG-gated gel-holes in yNup100FG particles and its surface. Tau values for *X. laevis* oocyte NPCs were calculated from (Sakiyama et al., 2016) (n = 3). Due to differences in their inner diameters, Tau was averaged over selected radii of ≦15 nm for yeast -CP NPCs, 20 nm for -CP Xenopus NPCs and 25 nm for the gel-holes, respectively. For the hydrogel surfaces, average tau values were calculated beyond radii of 25 nm (outside of gel-holes). Data are mean ± s.e.m. **b,** Molecular mechanism of the NPC permeability barrier. Left: When the CP is absent, the FG domains (green) function as a radial polymer brush whose dynamic movements fill the CT without forming a crosslinked meshwork. These carve out transient voids that shape-shift continuously. FG domain dynamics is slower at the scaffold anchor sites (gray) and faster along the pore axis. Right: During transport, the FG domains interact together with TFs and their cargoes to form a highly dynamic CP. This enhances the selective nature of the permeability barrier in the following ways: I. The CP reduces free aqueous space in the CT. II. Peripheral FG domains are more extended due to their interaction with TFs. III. The shearing (bristling) motion of the FG domains imparts an additional resistance against non-specific macromolecules. IV. Voids trap non-specific macromolecules. See Discussion for details.

Evidently, dynamicity is a highly conserved feature in NPCs from different organisms despite large differences in scaffold structure, FG number, mass, and concentration. On this note, there has been some confusion as to the estimated FG repeat concentration in the NPC, which varies from 50 mM (Frey and Gorlich, 2007), to 200 mg/ml (Labokha *et al*., 2013) and 6 mM (Schmidt and Gorlich, 2015). For clarity, we calculated the FG repeat and corresponding FG domain concentrations in isolated yeast NPCs (WT, ΔFG and Nsp1FGx2) and *X. laevis* NPCs based on a volumetric analysis of their CTs in both structures (Akey *et al*., 2022; Eibauer *et al*., 2015; Kim *et al*., 2018). These values are listed explicitly under “FG repeats” and “FG domains” and have been interconverted between different units (i.e., mg/ml and mM) in **Table S2**. Accordingly, we estimate that the FG domain concentration in WT NPCs is ∼96 mg/ml (∼2.3 mM), which is equivalent to an FG repeat concentration of ∼13 mg/ml (∼53 mM). These estimates are considerably lower than the FG domain concentration in hydrogels that show NPC-like selectivity, which was reported to be 200 mg/ml (3 - 4 mM; or 100-175 mM of FG repeats) (Schmidt and Gorlich, 2015).

## Discussion

### A dynamic NPC permeability barrier is greater than the sum of its parts

Our study provides insight into the molecular mechanism of NPC selective permeability across spatial, temporal, mechanical and functional levels. *In vivo*, NPCs always contain a large quantity of TFs (Kim *et al*., 2018), which makes up a significant portion of the CP and induces a dynamic “Kap-centric” organization of radial FG domain brushes along the pore axis *in situ* (**Figs. 1f and 3g; Fig. S8**). Hence, the selective nature of the permeability barrier is enhanced by a dynamic organization of the FG domains together with a preponderance of TFs that reduces nucleocytoplasmic leakage by dynamically outcompeting non-specific macromolecules (Kalita *et al*., 2022; Kapinos *et al*., 2017).

We emphasize that the dynamic molecular mechanism of the NPC permeability barrier is altogether more effective than the sum of its parts (**Fig. 6b**). First, the permeability barrier is stimuli-responsive such that the CP expands its vertical range by several nanometers in comparison to -CP WT NPCs (**Fig. 1i and j**). Second, the permeability barrier in +CP WT NPCs imparts a larger mechanoresistance than -CP WT NPCs. Third, FG domain fluctuations can impart both steric repulsive forces (i.e., resistance against compression) and lateral shear forces (i.e., resistance against sliding) (Klein, 1996) on non-specific cargoes that diffuse through the CT. Fourth, the shape-shifting voids trap non-specific entities (Winogradoff *et al*., 2022) thereby enforcing a more restrictive passive size limit on entry into the NPC and how deeply non-specific entities can traverse into the CT (**Fig. 1f-h** **and Fig. S4**). Fifth, the CP amplifies dynamic activity at the permeability barrier which is required to facilitate a rapid malleable response to local changes in the CT (**Fig. 1k and l**) such as dilation (see later). In this manner, we find that the CP represents a functional feature of the NPC permeability barrier. These features are fully consistent with polymer brush and virtual gating models for the state of the selective barrier (Kapinos *et al*., 2014; Lim *et al*., 2007; Lim *et al*., 2006; Rout et al., 2003; Rout *et al*., 2000; Schoch *et al*., 2012; Wagner *et al*., 2015) but provide a greater degree of granularity to our description of a dynamic “Kap-centric” organization.

### Confined polymer brushes underpin the dynamic nature of the NPC permeability barrier

Although FG domains sometimes show a predisposition to undergo LLPS *in vivo* and in purified forms, our results show that this behavior is actively suppressed at NPCs *in vivo*. LLPSs possess two distinct characteristics that are absent in NPCs, namely the untethered nature of the polymers that constitute LLPSs and their definition by self-organized surface tension (Feric et al., 2016; Taylor et al., 2016; Yamazaki et al., 2022). In the case of the NPC, this inability to phase separate appears to be imposed by the tethering of the FG domains to the scaffold at the walls of the central channel (**Fig. 5**) that constrains their ability to mix within the CT. Hence, the tethers are an essential contextual feature that should not be overlooked in *in vitro* studies, because they define the structural boundary of the translocation pathway and heavily modulate FG domain behavior.

Nevertheless, the proximity between anchor sites does influence how far the FG domains radially project into the CT. Surveying across organisms, FG domain lengths in endogenous FG Nups rarely exceed 700 amino acids (e.g., Human, Xenopus frogs, both budding and fission yeast, and the plant Arabidopsis (https://disprot.org/) (**Fig. S12**).

Moreover, when arranged by their approximate position along the CT’s vertical axis, those FG repeats found at the equator where the CT is narrowest are on average shorter than those found at the CT’s cytoplasmic and nuclear peripheries (Kim *et al*., 2018; Yamada *et al*., 2010) (**Table S1**). Based on their tethering constraints (**Fig. S9**), this suggests that the predicted average reach of these anchored FG regions and position of the anchor sites on the NPC’s scaffold is optimized so that deleterious entanglement is minimized *in vivo* (**Fig. 3**).

In another work (Otto *et al*., 2023), we show that the FG repeat domain of Nsp1 alters the phase state and reduces the aggregation of other FG Nups and this may contribute to maintaining the FG domains in the correct non-condensed state. Hence, we think that Nsp1 and potentially other FxFG domains are essential for optimizing FG domain dynamics in WT NPCs (**Fig. 1f and g**). Indeed, Nsp1, TFs and molecular chaperones (MLF2 and DNAJB6) have the ability to prevent the FG Nups from transitioning into dysfunctional condensates or aggregates (Kuiper *et al*., 2022; Prophet *et al*., 2022). Co-translational assembly and cytosolic chaperoning by Nsp1 (Otto *et al*., 2023), TFs (Hampoelz et al., 2019; Lautier et al., 2021; Lusk et al., 2002; Ryan et al., 2007; Seidel et al., 2022; Walther et al., 2003) and classical chaperones (Kuiper *et al*., 2022; Prophet *et al*., 2022) are emerging as a mechanism to keep FG Nups in an assembly or transport competent state. Indeed, TF-FG Nup co-translational assemblies may even constitute functional CPs that are crucial for preventing leakage across the NE during NPC assembly. In any case, we suggest that the formation of FG Nup condensates *in vivo* is a temporary phenomenon and related to regulation of NPC assembly (Colombi et al., 2013; Hampoelz *et al*., 2019; Makio et al., 2013).

### Shape-shifting voids within the passive permeability barrier

The presence of shape-shifting voids is also consistent with simulations which predict that passive cargoes diffuse through the permeability barrier in a time-dependent, percolation-like manner (Moussavi-Baygi and Mofrad, 2016; Winogradoff *et al*., 2022).

Furthermore, the circular arrangement of voids around the CP bears a striking resemblance to the location of low density regions at the periphery of the CT in cryo-EM structures that are accounted for by gaps between the anchor sites for bundles of FG domains (Akey *et al*., 2022; Kim *et al*., 2018) (**Fig. 6b**), as well as the Kapβ1 and transportin translocation pathways that are located at the periphery of the central channel (Chowdhury et al., 2022). The presence of these voids and their dynamic behavior may underlie the existence of spatially separated diffusion pathways in the NPC (Strawn *et al*., 2004).

### FG domain dynamics in dilatable NPCs

The NPC is a mechanically coupled system that can dilate, which increases the NPC’s circumference by 38% (Akey *et al*., 2022; Elosegui-Artola et al., 2017; Zimmerli *et al*., 2021). This theoretically increases the next-neighbor distance between anchor sites to 7.3 nm in the cytoplasmic lobe, 4.8 nm at the central waist and 10.1 nm in the nucleoplasmic lobe. Surprisingly, these distances are still sufficient so as to satisfy the brush requirement (Yamada *et al*., 2010) (**Fig. S9**). Still, a radial expansion of the spoke belt might further reduce the frequency of inter-FG Nup contacts, decreasing crowding, and so resulting in increased dynamics. We hypothesize that such increases in space and availability of free FG repeats could even potentially lead to an increase in the number of TF molecules within the CT to maintain a steady state concentration that is comparable to non-dilated NPCs. In this way, the balance of FG density and TF concentration in the CT may help buffer the properties of the selective barrier during NPC diameter changes.

### Altering the balance of FG content in the CT abrogates normal NPC transport function

As well as the correct TF content, correct organization of the amount, type and position of different types of FG repeat domains in the CT appears important for NPC function. Our mutant Nsp1FGx2 NPCs with increased FG domain mass show inherently slower dynamics (**Fig. 3g**) likely due to increased crowding in the CT, higher FG concentration, and the increased propensity for opposing FG Nups to interact. This resulted in an unnatural meshwork-like behavior that tightened the NPCs’ passive permeability against high molecular weight reporters and caused active transport defects *in vivo* (**Fig. 3h-j**), indicating that overly strong crosslinking interactions can be deleterious. This behavior is analogous to meshwork-forming proteins that inhibit nucleocytoplasmic transport by accumulating at NPCs, such as neuropathological amyotrophic lateral sclerosis (ALS)-associated poly-dipeptides (Shi et al., 2017) and phospho-Tau in Alzheimer’s disease (Eftekharzadeh et al., 2018). The same rationale holds true for wheat germ agglutinin (Finlay et al., 1987) and dominant-negative mutants of Kapβ1 (Kutay et al., 1997) that block NPCs *in vitro*. Hence, we intuit that TF enrichment at NPCs to form CPs, in conjunction with the dynamicity of the polymer brush-like FG domains, provides a reversible means of tuning the passive permeability barrier versus optimizing selective transport to form an exquisitely-tuned but robust selectively permeable gate. Notably, this intricate balance is lost in Nsp1FGx2 NPCs, where meshwork formation artificially overtightens the permeability barrier, which impairs selective transport *in vivo*.

### Condensates do not represent the bona fide state of FG domains in NPCs

Biomolecular condensates can undergo a liquid-to-solid transition to form heterogenous aggregates (Ahmad et al., 2022). Prominent examples include ALS-related proteins FUS (Patel et al., 2015) and TDP-43 (Johnson et al., 2009). Likewise, we find that FG hydrogels are structurally heterogenous assemblies composed of granular aggregates, amyloid fibrils, and gel-holes that pockmark the surface and permeate the bulk in a manner reminiscent of “Swiss cheese” (**Fig. 4c-e**). Our data indicate that such condensates do not represent the *bona fide* state of FG domains in NPCs and that condensate formation does not occur in healthy NPCs; however, sufficient alterations of FG domains at the NPC may overcome this suppression and lead to malfunctioning disease states. This is in agreement with studies showing that FG Nups form dynamic and transient storage receptacles or assembly intermediates in a liquid-like state by the action of TFs and Nsp1 (Hampoelz *et al*., 2019; Kuiper *et al*., 2022; Lautier *et al*., 2021; Lusk *et al*., 2002; Otto *et al*., 2023; Prophet *et al*., 2022; Ryan *et al*., 2007; Walther *et al*., 2003).

To be precise, our *in situ-*observed “Kap-centric” organization deviates considerably from models that invoke biomolecular condensation-type behavior such as the selective phase model, which likens the NPC permeability barrier to an *in vitro* FG hydrogel (Frey and Gorlich, 2007; 2009; Frey *et al*., 2006; Hülsmann *et al*., 2012; Labokha *et al*., 2013; Schmidt and Gorlich, 2015). Several lines of evidence contradict the view that the FG domains alone undergo extensive crosslinking to cohere into a “saturated hydrogel” in WT NPCs. We call particular attention to: (i) the continuous dynamic motion of the FG domains; (ii) an insufficient extensibility of the FG domains to reach each other across the CT; (iii) the dynamicity of shape-shifting voids that emphasizes poor cohesion between the FG domains; (iv) the lack of fusion between Nup100FG LLPS condensates and NPCs *in vivo*; and (v) the fact that actual FG domain-meshwork formation in Nsp1FGx2 mutant NPCs disrupts nucleocytoplasmic transport *in vivo*.

Still, how do hydrogels that do not bear any resemblance in size, shape or form to NPCs display NPC-like characteristics (Frey and Gorlich, 2007; 2009; Frey *et al*., 2006; Hülsmann *et al*., 2012; Labokha *et al*., 2013; Schmidt and Gorlich, 2015)? An explanation for why FG hydrogels mimic some NPC behaviors lies in the presence of FG repeat-lined gel-holes that superficially mimic the NPC. Curiously, the mean of the gel-hole diameters is ∼50 nm (**Fig. 4g**), which is similar to the CT diameter. Importantly, this agrees with the prior finding that FG Nup hydrogels contain numerous NPC-sized nanopores that are coincident with large networks of amyloid fibers that maintain the ability to bind TFs (Milles *et al*., 2013). Other pore-like conformations have also been detected in TDP-43 aggregates (Johnson *et al*., 2009). These gel-holes clearly do not correspond to the much smaller “gaps” that have been hypothesized to lie between hydrogel meshes in the selective phase model (Frey and Gorlich, 2007; 2009; Frey *et al*., 2006; Hülsmann *et al*., 2012; Labokha *et al*., 2013; Schmidt and Gorlich, 2015). Rather, we surmise that the gel-holes crudely mimic the lined polymer brush of the NPC CT, acting as selectively gated passageways into the particle interior. Collectively, the results presented here and those of others (Milles *et al*., 2013) indicate that utmost care should be exercised when drawing inference from the bulk behavior of *in vitro* reconstituted FG domain assemblies to describe nanoscopic NPC function. Moving forward, we suggest that the necessarily high degree of organization in the NPC may be better replicated *in vitro* using biomimetic NPC-like nanopores (Fisher et al., 2018; Fragasso et al., 2022; Jovanovic-Talisman et al., 2009; Ketterer et al., 2018; Kowalczyk et al., 2011; Shen et al., 2023).

### Limitations of the study

Technical caveats: 1. HS-AFM data depends on the quality and dimensions of its tip – we only use pristine tips with less than 2 nm radius. 2. HS-AFM more easily visualizes FG Nups that are oriented in the XY plane than those (if any) that align in the Z axis. 3. HS-AFM cannot resolve features beyond the lowest accessible depths. Here, the most important features - the dynamic fluctuations and voids – are mostly resolved at depths closer to the midplane. 4. HS-AFM is a single molecule method that incurs experimental limits on sample size. We anticipate that improvement in both preparation methods and HS-AFM technologies (Ando, 2018) will allow further characterization of NPCs, including those in the process of discernable cargo transport.

## Acknowledgements

R.Y.H.L. is supported by the Schweizerischer Nationalfonds zur Förderung der Wissenschaftlichen Forschung (Swiss National Science Foundation; grant no. 310030_201062), the Biozentrum and the Swiss Nanoscience Institute. T.K. is supported by a Swiss Nanoscience Institute PhD Fellowship. J.F-M. acknowledges grant funding from National Science Foundation (NSF-1818129) and Spanish Ministry of Science and Innovation (PID2020-116404GB-I00; MCIN/AEI/10.13039/501100011033). M.D-I. is supported by the Investigo Program (Lanbide-Servicio Vasco de Empleo) funded by Next Generation EU (Plan de Recuperación, Transformación y Resiliencia). B.T.C., M.P.R., and A.S. are supported by NIH P41 GM109824. M.P.R. is supported by NIH R01 GM112108 and R01 GM117212. A.S. is supported by NIH R01 GM083960. PG and LV are supported by the Netherlands Organization of Scientific Research; grant no. OCENW.GROOT.2019.068 and VI.C.192.031. We thank A. Steen and T. Bergsma for providing advice and purified Nup100FG and E.C. Riquelme Barrientos for providing the plasmid mCherryWALP. We also acknowledge P. Upla and R.E. Marin as well as the Imaging and Bio-EM facilities at the Biozentrum for support. We also thank nanotools GmbH for permission to use the TEM image of the QUANTUM-AC10-SuperSharp HS-AFM tip shown in Fig. 1a.

## STAR METHODS

### KEY RESOURCES TABLE

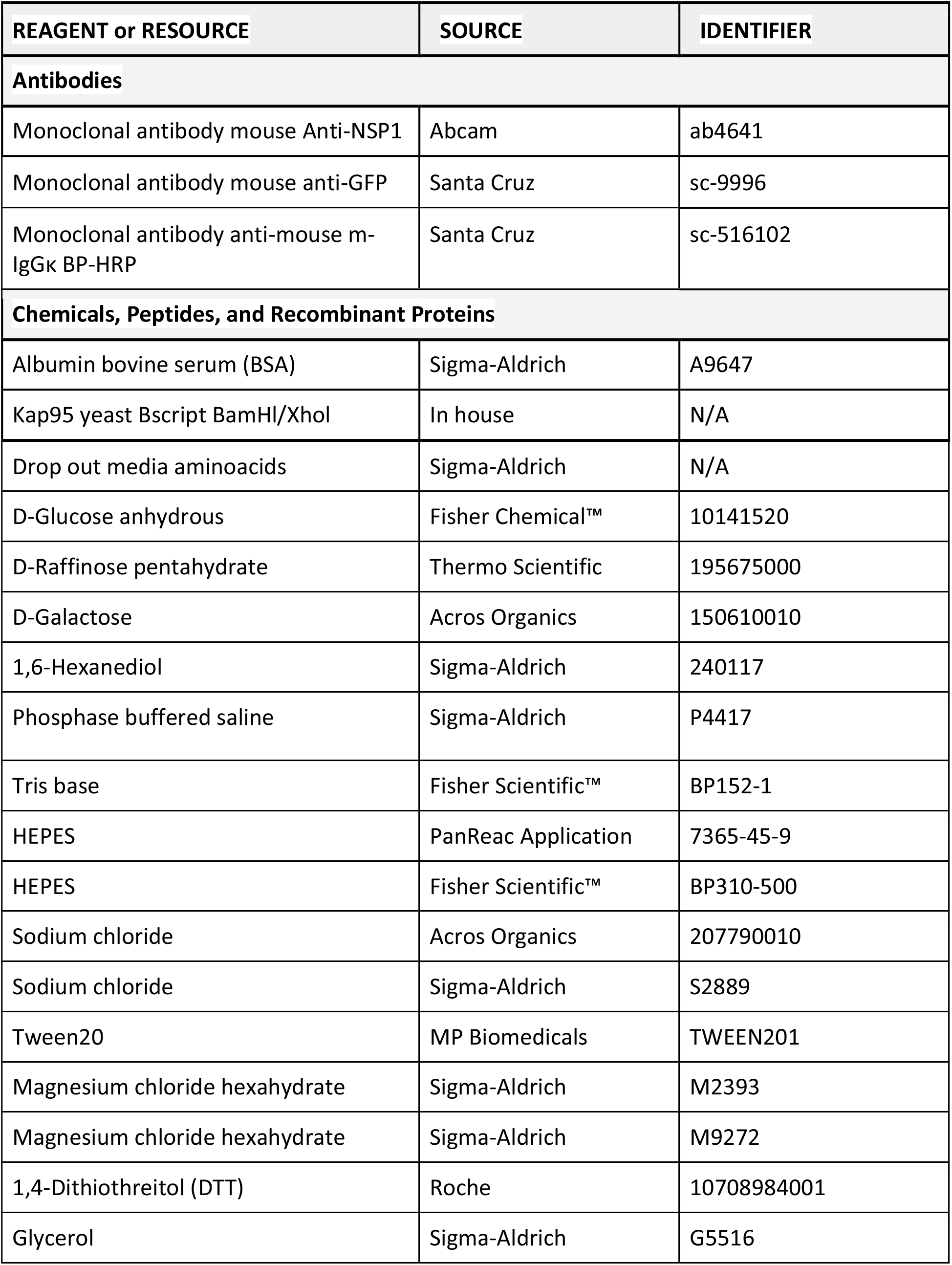
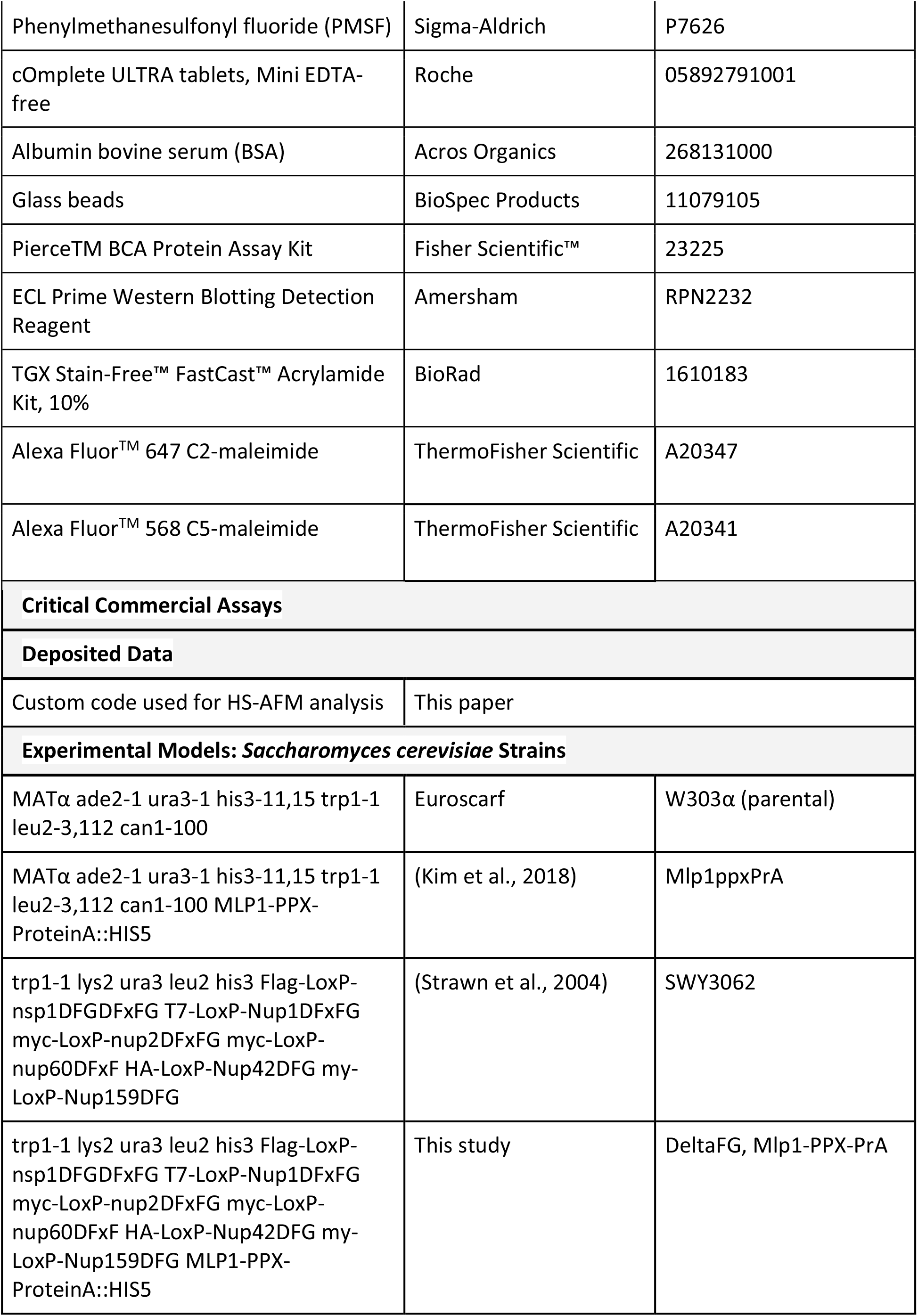
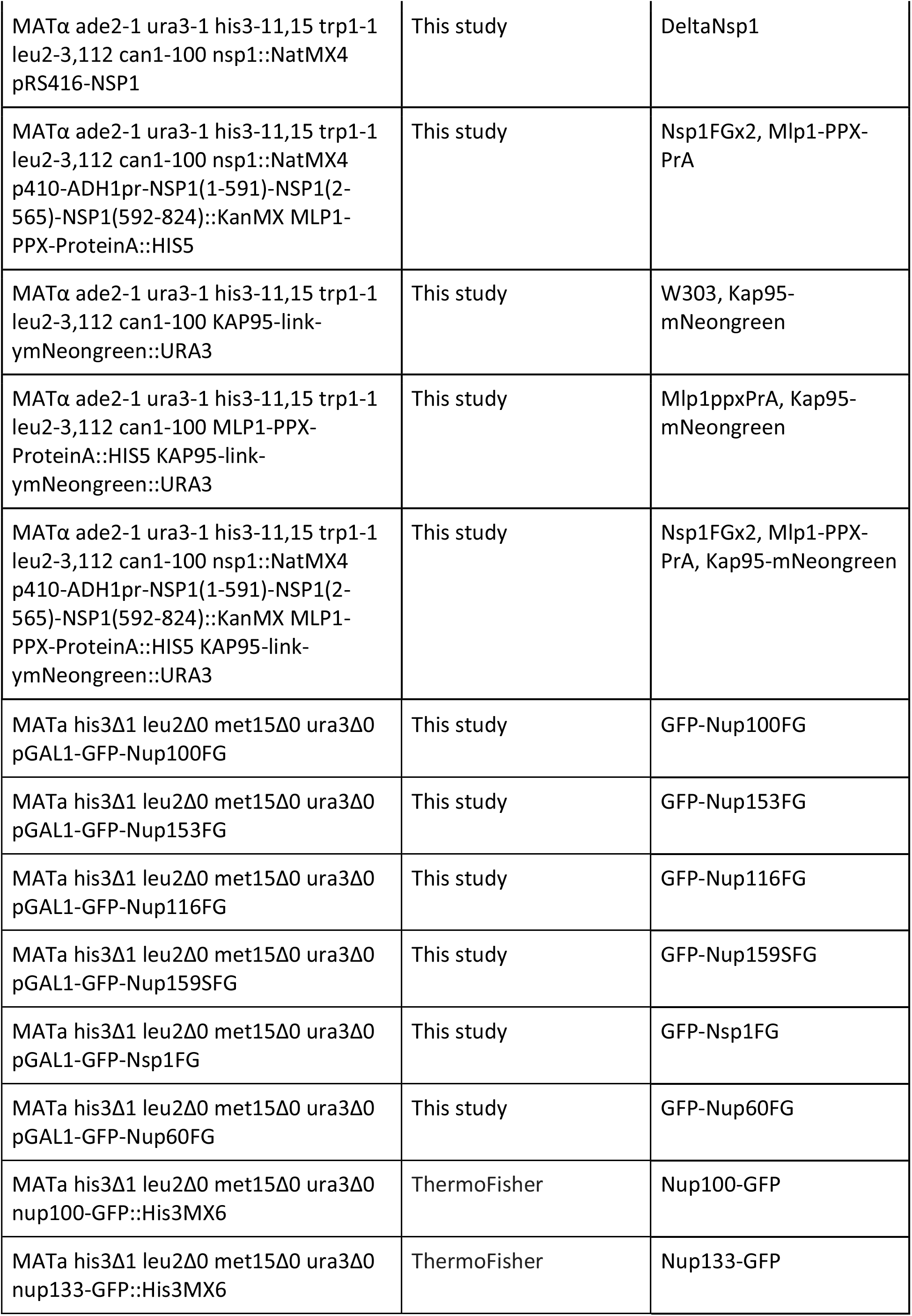
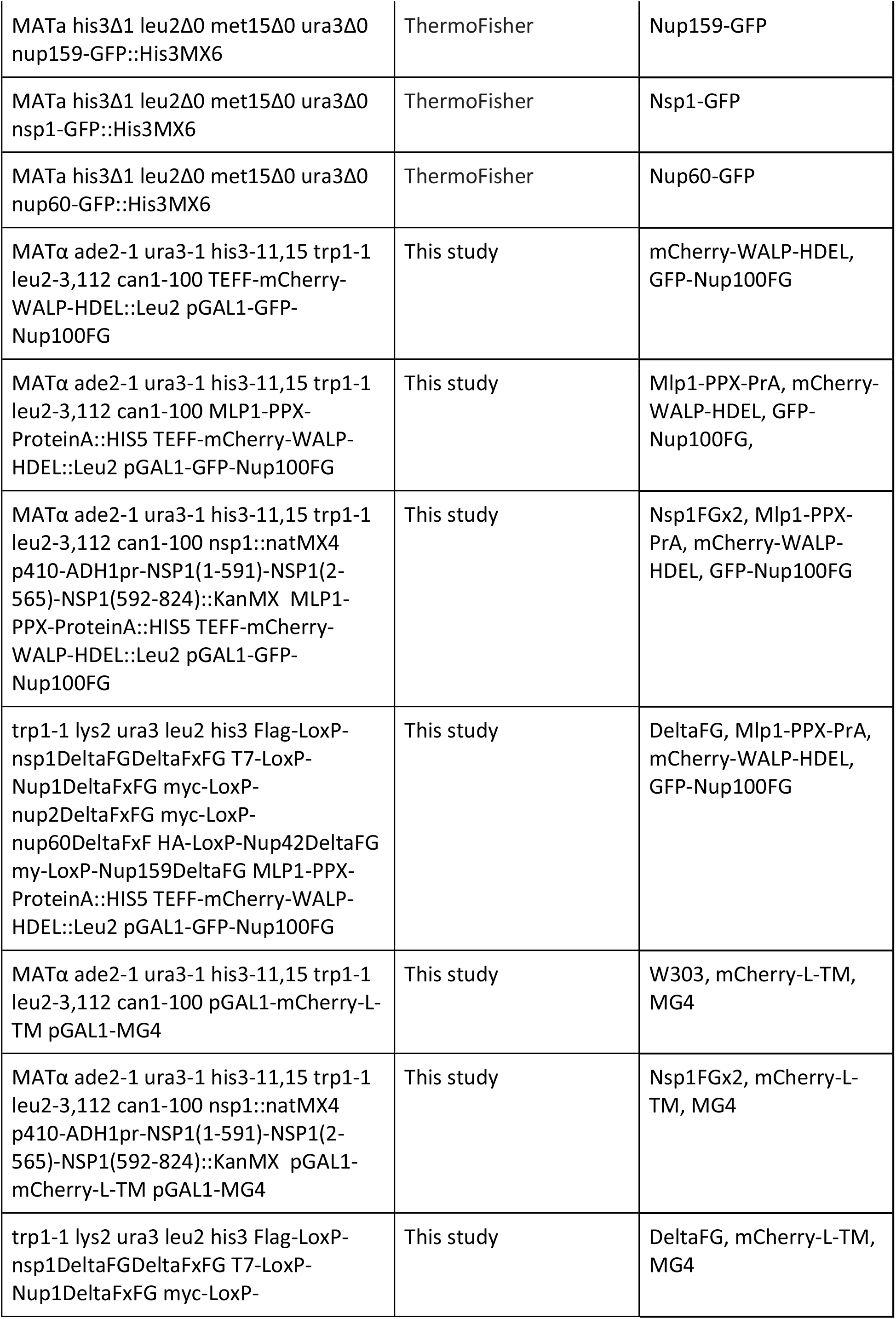
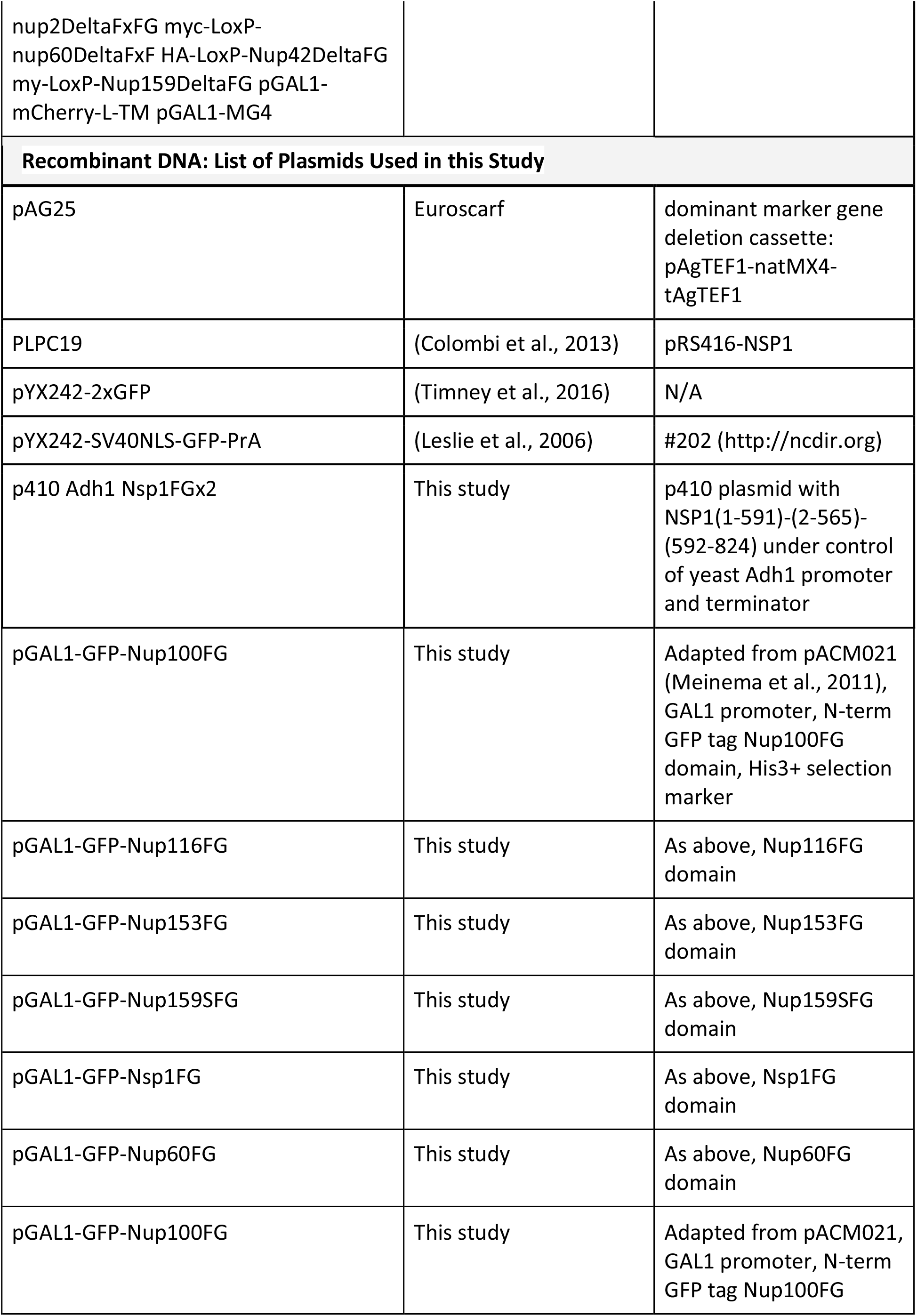
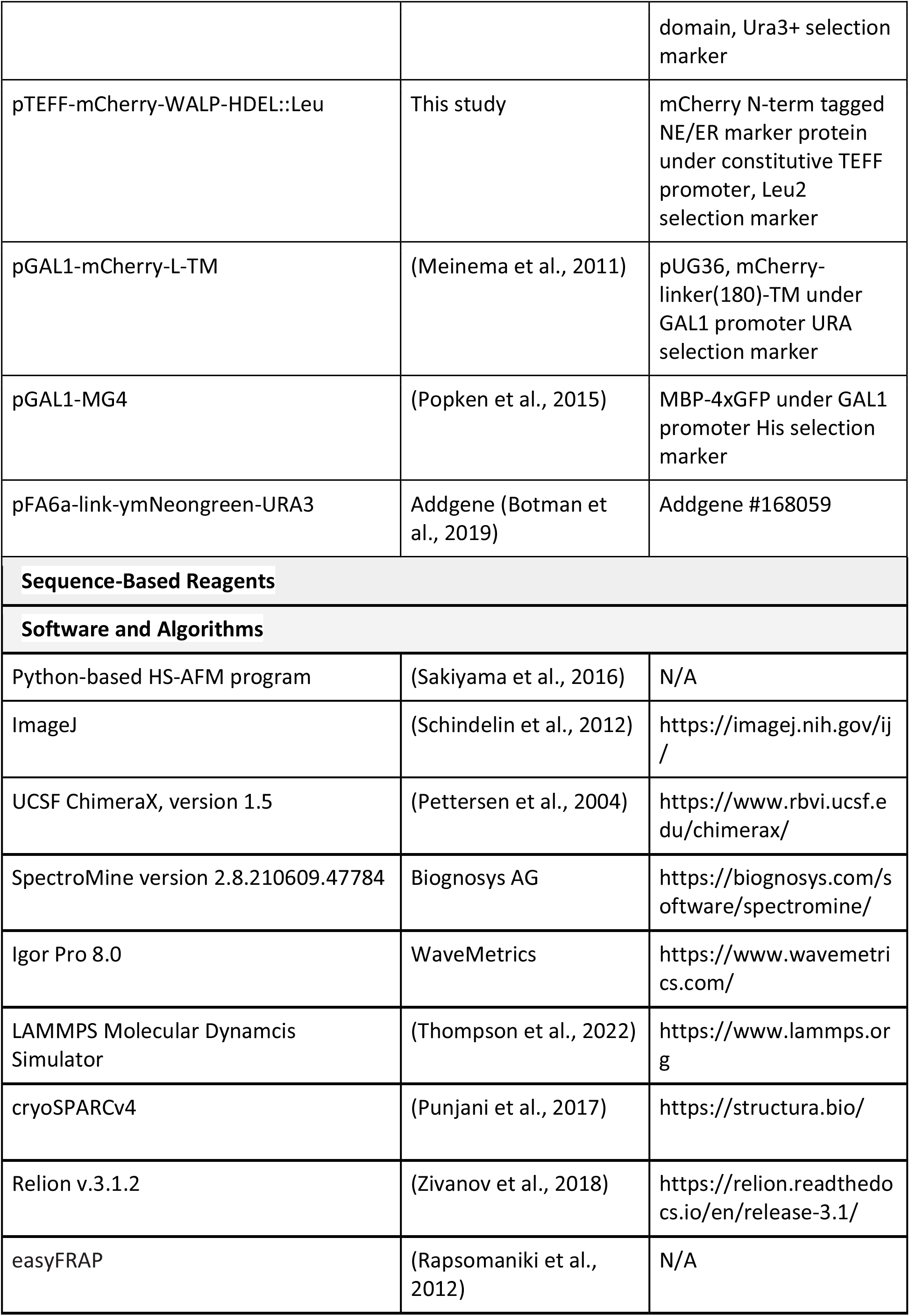
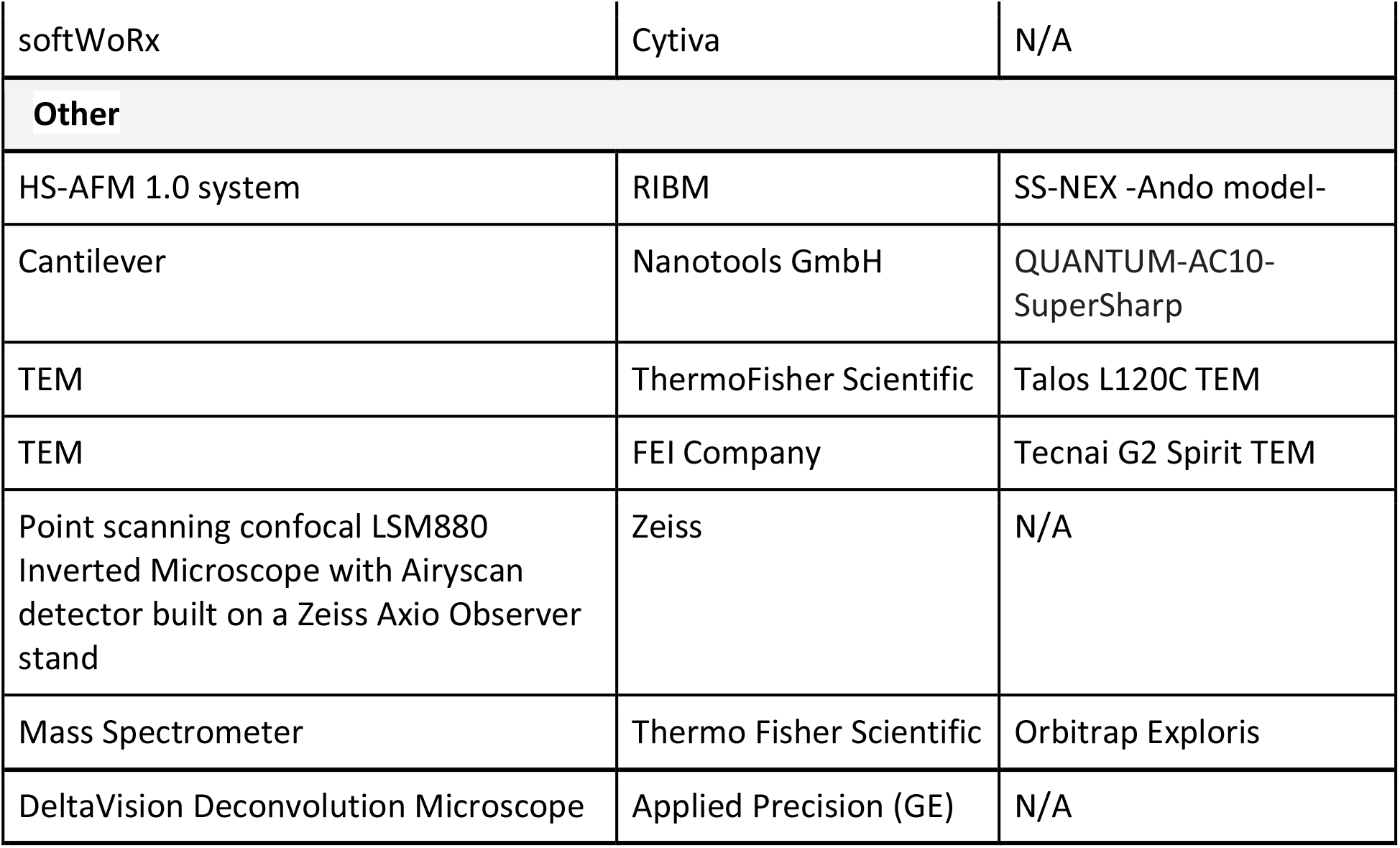

### RESOURCE AVAILABILITY

#### Lead contact

Further information and requests for resources and reagents should be directed to and will be fulfilled by the lead contacts, Michael Rout and Roderick Lim.

#### Materials availability

Yeast strains and plasmids generated in this study will be distributed without restriction upon request.

#### Data and code availability

HS-AFM data has been deposited in --.

Mass spectrometry data files have been deposited in --.

Raw image data files are available in the Zenodo public repository under the identifier --.

Software scripts and simulation data are available at https://github.com/ --.

Any additional information will be made available from the lead contact upon request.

### EXPERIMENTAL MODELS AND SUBJECT DETAILS

#### Yeast strains and materials

All *S. cerevisiae* strains used in this study are listed in the Key Resources Table. Unless otherwise stated, strains were grown at 30°C in YPD media (1% yeast extract, 2% bactopeptone, and 2% glucose) supplemented with adenine hemisulfate (40 mg/l; Sigma).

### METHOD DETAILS

#### Yeast strains construction

Yeast strains were constructed using standard molecular genetic methods (Longtine et al., 1998). For the construction of the Nsp1FGx2 mutant, the Nsp1 open reading frame was knocked out from a strain carrying plasmid PLPC19 by using a cassette amplified from plasmid pAG25 that introduces a nourseothricin resistance marker (DeltaNsp1) (Goldstein and McCusker, 1999). Using NSP1 from the S288C reference genome as a basis, a gene was constructed to effectively double the FG region of Nsp1 by repeating residues 2-565 between residues 591 and 592, resulting in the sequence: Nsp1(1-591)-(2-565)-(592-824) (Nsp1FGx2). In the Nsp1FGx2 gene, the NSP1 intron was removed and the gene was codon optimized and synthesized by Twist Bioscience. The Nsp1FGx2 gene was cloned into the p410 plasmid under control of the Adh1 promoter and terminator. DeltaNsp1 yeast were transformed with the p410-Adh1-Nsp1FGx2 and plated on YPD+G418. Single colonies were passaged three times on YPD+G418 at 30°C and then plated on SC-Ura and SC+5-FOA plates. Colonies that had naturally lost the PLPC19 plasmid (SC-Ura negative, SC+5-FOA positive) were chosen for downstream analysis. The sequences for all the oligonucleotides and vectors used in this study are available upon request.

#### Affinity-purification of endogenous S. cerevisiae NPCs

We used our previously published method for the isolation of endogenous whole NPCs from *S. cerevisiae* (Akey et al., 2022; Kim et al., 2018; Nudelman et al., 2022). Briefly, Mlp1 was genomically tagged in each strain with PrA preceded by the human rhinovirus 3C protease (ppx) target sequence (GLEVLFQGPS). Cells were grown in YPD media at 30°C until mid-log phase (∼3x10^7^ cells/ml), harvested, frozen in liquid nitrogen and cryogenically lysed in a planetary ball mill PM 100 (Retsch) (http://lab.rockefeller.edu/rout/protocols). Affinity purification was performed in resuspension buffer (20 mM HEPES/KOH pH 7.4, 50 mM potassium acetate, 20 mM NaCl, 2 mM MgCl_2_, 0.5% (w/v) Triton X-100, 0.1% (w/v) Tween-20, 1 mM DTT, 10% (v/v) glycerol, 1/500 (v/v) Protease Inhibitor Cocktail (Sigma)) and native elution by protease cleavage was achieved in a similar buffer without Triton X-100 or protease inhibitors. Protease inhibitors were added to the purified NPCs and either conserved at 4°C or frozen in liquid nitrogen and stored at -80°C.

Successful incorporation of Nsp1FGx2 was determined by SDS-PAGE analysis and Western blotting of purified NPCS as follows. NPCs from Nsp1FGx2, Mlp1-PPX-PrA cells grown and lysed as described above were purified using a citrate resuspension buffer (20 mM HEPES, pH 7.4, 150 mM NaCl, 250 mM sodium citrate, 1% v/v Triton X-100, 1x protease inhibitor cocktail (PIC)) as described in (Hakhverdyan et al., 2021). Affinity captured NPCs were eluted from beads by addition of 20 μl of 1x LDS (lithium dodecyl sulfate) loading buffer (Thermo Fisher) and vortexing for 10 minutes at room temperature. Eluted NPCs were run on an NuPAGE 4-12% gel for 60 minutes and analyzed by Coomaisse or western blot. For western blots, NPCs were transferred onto PVDF membranes overnight at 4°C, blocked with 5% powdered fat free milk in TBST (0.1% Tween20) and probed overnight with primary antibody monoclonal mouse anti-Nsp1 (abcam) (1:5000). Membranes were washed with TBST, reprobed with secondary antibody anti-mouse (1:5000) for 1 hour at room temperature, then treated with ECL prior to imaging with ImageQuant 4000 LAS.

#### Quantitative mass spectrometry analysis of affinity purified NPCs

The affinity captured NPCs (5 μg per sample) were concentrated by pelleting at 40,000 r.p.m. for 20 min at 4°C in a TLA 55 rotor (Beckman). Pelleted NPCs were solubilized by addition of a final 1x of NuPAGE LDS Sample Buffer (Invitrogen) and 50mM DTT, and incubation at 72°C for 10 min. Samples were then cooled down to room temperature and treated with a final 30 mM of iodoacetamide (Sigma), at room temperature in the dark for 30 min. Samples were then loaded into 4% (37.5:1) stacking acrylamide SDS-PAGE gels prepared in-house. The resulting stacked bands (gel plugs) were stained by coomassie, excised and processed for quantitative mass spectrometry analyses.

Proteins in gel plugs were digested and peptides were extracted as described in (Bosch et al., 2021). The purified peptides were bound to C18 StageTips. Peptides eluted from the StageTips were analyzed by LCMS Easy-nLC system coupled to a Thermo Orbitrap Exploris mass spectrometer (Thermo Fisher Scientific). SpectroMine (version 2.8.210609.47784, Biognosys AG) software was used to identify and quantitate the mass spectrometric data by means of label-free quantitation (LFQ) (Bosch et al., 2021). The protein LFQ outputs from SpectroMine were further analyzed with Microsoft Excel. The absolute stoichiometries of the proteins were determined by normalizing the summed copies of Nup192, Nup188, Nup170, Nup157, Pom152 and Nic96 per NPC to 112 copies (i.e., 16 for Nup192, Nup188, Nup170, Nup157 and Pom152, and 32 for Nic96).

#### Fitness and growth assays

To analyze their fitness at different temperatures, strains were grown in liquid yeast extract peptone dextrose (YEPD) media overnight at 30°C, and cells were counted and diluted to a final concentration of 20,000 cells/ml. Four 10-fold serial dilutions were made and spotted on YEPD plates that were incubated at 25, 30, and 37°C for 1–2 d. Three biological replicas of each experiment were performed. A ΔNsp1 + Nsp1 strain was also analyzed to ensure no deleterious effect of the gene deletion background (not shown). Plates were imaged using a Versadoc imaging system (linear detection range; Biorad). Semiquantitative estimation of fitness was performed using ImageJ (National Institutes of Health) as previously described (Fernandez-Martinez et al., 2012).

For growth curves, strains were grown in biological replicas as described above, inoculated in 96-well plates (150 µl) and grown with 19 hertz agitation at each temperature for 30-50 hours in a Biotek Synergy HT microplate reader. OD600 reads were calibrated (Small et al., 2020) and transformed into cell counts for plotting the curves.

#### Recombinant protein expression and purification

Yeast Kap95 was recombinantly purified in-house from *E. coli* BL21DE3. The protein was purified from a Ni-NTA column. The protocol is as follows: 3 liters of BL21DE3 bacteria were induced at OD_600_ = 0.6 with 0.5 mM IPTG at 21°C overnight. After harvesting bacteria, lysis buffer in the following composition (10 mM Tris pH 7.5, 100 mM NaCl, 1 mM DTT, 10 mM Imidazole +DNase, Lysozyme, Pefabloc, PIC Complete) was used and incubated for 1 hour at 4°C. Lysate was sonicated by a stab-sonicator at 30% amplitude, 3 sec pulse and 4 sec off, for a total of 5 min pulse. The supernatant from the lysate was separated from unlysed debris after a spin at 25000 g, for 1 hour at 4°C. The supernatant was loaded onto a 5 ml Ni-NTA column (GE Healthcare). After flowthrough and wash steps, the proteins were eluted in a lysis buffer supplemented with 500 mM imidazole.

#### High-speed atomic force microscopy

All HS-AFM movies in this study were acquired using a HS-AFM 1.0 system (RIBM, Japan) with a standard scanner operating in tapping mode at a maximum scan speed of 150 ms per frame. Pristine QUANTUM-AC10-SuperSharp cantilevers (nanotools GmbH) bearing a tip radius of ≤ 2 nm (**Fig. 1a**), nominal spring constant of 0.1 N/m, resonant frequency of ∼0.5 MHz, and a quality factor of ∼2 in buffer were used in all experiments. The typical set point amplitude *A_set_* was 80-90% of the free cantilever oscillation amplitude *A_free_*, which was set to 2-3 nm (Ando, 2018; Uchihashi et al., 2012). 3 µl droplet containing isolated NPCs were dispensed on freshly cleaved mica surfaces for 3 min at RT. Any excess was removed by rinsing the sample with buffer (20 mM HEPES-KOH pH 7.4, 20 mM NaCl, 2 mM MgCl_2_). As an exception, isolated ΔFG NPCs were immobilized on 3-aminopropyltriethoxy silane-functionalized mica (APTES-mica) which was prepared by incubating a droplet (3 µl) of 0.1% APTES on a freshly cleaved mica surface for 5 min followed by thorough rinsing with pure water.

#### Negative stain transmission electron microscopy and image processing

For negative stain analysis of purified yeast NPCs, 5 µl of purified sample was adsorbed onto 400 mesh carbon-coated grids which were rendered hydrophilic using a glow-discharger at low vacuum conditions. Grids were washed with 3 drops of water or TEM buffer (20 mM HEPES, 20 mM NaCl, 1 mM DTT) and subsequently stained with 5 µl drops of 2% (w/v) uranyl acetate. Samples were examined either with a Tecnai G2 Spirit TEM (FEI Company, USA) operating at 80 kV accelerating voltage or a Talos L120C TEM (ThermoFisher Scientific, USA) operating at 120 kV. On the Tecnai G2 Spirit, TEM images were recorded with an Olympus Veleta camera 4k using EMSIS Radius software at nominal magnification of 49kx. On the Talos L120C TEM, images were recorded with a Ceta 16M Pixel CMOS camera using the TIA software at a nominal magnification of 28kx. The software packages cryoSPARCv4 (Punjani et al., 2017) and RELION v.3.1.2 (Zivanov et al., 2018) were used for particle picking and 2D classification.

#### Brownian dynamics simulation of the NPC

We represent the yeast NPC as a coarse-grained (CG) model consisting of three main components: 1) scaffold Nups consisting comprising the cytoplasmic, nuclear and inner rings, 2) the flexible FG Nup domains and 3) the nuclear envelope (NE). The scaffold is represented as a fixed set of spherical CG beads derived from the previously published model of the yeast NPC (Kim et al., 2018). The FG Nup domains are represented as a chain of CG beads where each bead corresponds to 20 residues and is represented as a sphere with radius of 6 Å. The positions of the anchor points for each FG Nup are also derived from the Kim 2018 model. Each CG bead has a single binding patch represented as a sphere with radius of 1 Å. The NE is represented as a cylindrical pore of radius 300 nm embedded in two parallel planes set at a distance of 38 nm apart.

All non-bonded interactions in the system are modeled using a 12-6 Lennard-Jones interaction potential. The bonded interactions are modeled as a Finite-extensible non-linear elastic (FENE) potential. Binding patch particles and the parent CG bead are treated as a rigid unit. Interactions between all particles and the NE membrane are modeled as a harmonic repulsive term that prevents particles from crossing the NE surface into the NE lumen.

We simulate the dynamics of the FG Nup domains using the LAMMPS molecular dynamics simulation package (Thompson et al., 2022) with the standard Langevin dynamics algorithm at a temperature of 297K, a damping coefficient of 700 fs and a timestep of 200 fs. The particles in the NPC scaffold and the position of the NE membrane surface are fixed in place during the simulation. We constrain all rigid bodies using the rigid body constraints package in LAMMPS. The initial positions of the scaffold and FG-nups are taken from the Kim2018 model files. We then perform a steepest descent minimization algorithm to remove steric clashes followed by an equilibration run. We increased the temperature of the simulation gradually over the equilibration run over a total of 0.1 μs. Finally, we simulate 1.44 × 10^8^ timesteps of 500 fs for a total of 72 μs of simulation time.

#### Simulating a high-speed atomic force microscope experiment

Simulated HS-AFM images are computed from a single molecular trajectory through the following rasterization scheme. First, we define the imaging plane as a 40 x 40 rectangular grid of pixels centered on the origin and aligned with the XY-plane at a height of Z = 0 (i.e., equator of the NPC channel). Each pixel is defined as a square cell of size 0.9875 nm, which corresponds to our typical HS-AFM imaging condition. The value of each pixel is assumed to be proportional to the total force required to compress all particles in the simulation located above the imaging plane (z > 0) and within the field of the AFM imaging tip (modeled as a cylinder of radius 3 nm centered over the given pixel) vertically to the imaging plane. The force required to compress the particles to the imaging plane is modeled as a linear response function. The HS-AFM image is calculated starting at (1, 1) position on the grid with fast scanning along the x-axis, i.e. row-wise. To mimic the HS-AFM acquisition scheme, a single pixel is calculated from one CG molecular dynamics trajectory snapshot.

#### HS-AFM data processing

All HS-AFM movies were corrected for drift by an in-house Python-based software that also converted the file into tiff format. HS-AFM movies were analyzed in ImageJ and self-written analysis routines in Python. Image filtering, contrast adjustment, height/diameter measurement and area calculation were performed by ImageJ. Briefly, the HS-AFM frames were shifted to a reference frame by minimizing their mean square difference to correct for Z-drift. A flattening filter with first order polynomial plane was applied to compensate for XY-tilting. The frames were then corrected for XY-drift based on a manually selected reference frame using the previously used image registration algorithm (Sakiyama et al., 2016). The frames were cropped to ROI into a square afterwards. ACF analyses was performed without denoising filters except the flattening filter unless stated otherwise. Otherwise, displayed HS-AFM images are treated by a 2D Gaussian filter with a 1-pixel standard deviation followed by bicubic interpolation with a scale factor of 2 in the x and y direction. The HS-AFM analysis workflow is outlined in **Fig. S5**.

#### Alignment of zoom-in to zoom-out images

To set a pore depth relative to its NPC rim, raw zoom-in images of the CT and the zoom-out images of entire NPCs were first averaged. The average zoom-in image was then aligned with the average zoom-out image in the XY and Z directions by rescaling the pixel size to match the pixel size, identifying a maximum normalized cross-correlation coefficient and minimizing the mean square difference between the two images. The Z-shifted value obtained by the alignment was added to pixels in all zoom-in images followed by subtraction of the average maximum height, which was calculated from each of the 8 spokes of the NPC by searching an area of 15 x 15 pixels (implemented in ImageJ) in the average zoom-out image.

#### Average HS-AFM image of the NPC

We manually selected image stacks of NPCs scanned with a pixel size of 1 nm/pixel. After the drift corrections, the raw images in each stack were averaged over a corresponding pixel position. Then, the average image was rotated and aligned by phase cross-correlation. The rotational alignment was performed twice for accuracy. Subsequently, all images were averaged over.

#### Height fluctuation and average Tau representations

To plot peak-peak fluctuations in the Z-axis, height fluctuations taken from all kymographs were plotted in normalized height heatmaps as a function of distance from center. For illustration of average Tau values as a function of distance from center, the Tau values estimated from each HS-AFM movie were averaged over 1-nm intervals of distance from center. The averaged values were then plotted in polar coordinates displaying a 360-fold symmetrized heatmap as a function of distance from center.

#### HS-AFM noise baseline analysis

The HS-AFM noise baseline was determined by measuring the height fluctuations on mica surfaces and lipid membranes with hole-like defects, sampled with the same imaging conditions as NPCs, which gave RMS amplitudes of 0.13 nm and 0.35 nm, respectively. Slightly larger RMS amplitude on lipids is due to diffusion of lipid molecules. When compared with the RMS amplitude (2.8 nm) of FG Nups, RMS amplitudes on mica and lipids are significantly smaller. This eliminates the possibility that height fluctuation in CT is due to intrinsic HS-AFM noise (**Fig. S7**).

#### HS-AFM void analysis

To facilitate measuring void area frame by frame in ImageJ, a 2D Gaussian filter with a 1-pixel standard deviation followed by bicubic interpolation with a scale factor of 2 in the x and y directions were applied to all HS-AFM movies. The area was then measured in ImageJ by identifying the pixel values that fell below a user-defined threshold and by changing the threshold with 2 nm increments.

#### Calculation of next-neighbor distance for WT and ΔFG NPCs

We extracted the anchor point coordinates for the FG repeat domain from the integrative localization map of the yeast NPC, which has a precision of 9 Å (Kim et al., 2018). For the wildtype analysis, we included the anchor sites of Nup1.601-636 (i.e., residues 601-636 of Nup1), Nup2.301-350, Nup49.201-269, Nup57.201-286, Nup60.351-398, Nup100.551-575, Nup116.751-775, Nup145.201-225, and Nup159.1082-1116; we omitted the anchor site of Nup42 from the analysis because it is likely to fluctuate spatially. Additionally, the anchor sites of Nsp1, Nup159, Nup1, and Nup60 were omitted in the deletion mutant analysis. For a single anchor site, the nearest-neighbor distance is the minimum Euclidean distance to any other anchor site within a single spoke of the NPC, including other copies of the same anchor site.

#### In vivo nucleocytoplasmic transport reporter assays

Indicated yeast strains were transformed with 2 μl plasmid pYX242-SV40NLS-GFP-PrA expressing the SV40NLS-GFP-PrA reporter (Leslie et al., 2006) or 2xGFP (pYX242-2xGFP). Cells were grown at 30°C overnight in 5 mL of SC-Leu media (containing 1% (m/v) succinic acid, 0.6% (m/v) NaOH, 160% (m/v) dropout powder, 2% D-glucose and 1xYNB+AS (Yeast Nitrogen Base medium without amino acids supplemented with ammonium sulfate) supplemented with 1 µg/ml ampicillin and 0.2 mg/ml of adenine hemisulfate. After OD_600_ reached 0.6-1 the yeast cells were 1:20 diluted into the fresh SC-Leu medium and allowed to grow for another 4 h. Since Kap95-mNeonGreen strains were genomic integrations, in this case the regular 1xYPD medium (1% yeast extract, 2% peptone, 2% glucose; from Sigma) was used supplemented with the 0.25% Adenine Hemisulfate. Finally, yeast cell cultures were centrifuged at 500 rpm for 5 min and a fraction of the pelleted cells was re-suspended in a small amount of fresh medium and placed into 8-well ibidi slides, that had been prior treated with poly-lysine. After the yeast cells were settled, the ibidi slides were mounted on to the confocal microscope stage for fluorescent imaging.

Passive permeability for MG4 of the W303, ΔFG, and Nsp1FGx2 stains was determined as previously described (Popken et al., 2015). For the Δnsp1 p410-Nsp1FGx2:KanMX strain medium was supplemented with G418 (200 μg/ml). Prior to microscopy, 1 ml of cell yeast culture was centrifuged and a fraction of cells were resuspended and mounted on glass slides.

#### In vitro phase separation of yNup100FG hydrogels

Nup100 hydrogels were prepared following an established protocol (Schmidt and Gorlich, 2015). 100 µM yNup100FG was purified as described (Kuiper et al., 2022). For HS-AFM experiments, yNup100FG condensates were formed by first placing a 2.9 µl droplet of TBS buffer (150 mM NaCl and 50 mM Tris–HCl, pH 8) on freshly cleaved mica, and mixing with 0.1 µl of 100 µM yNup100FG. This resulted in a total volume of 3.0 μl with a final concentration of 3.3 µM yNup100FG. The samples were then gently rinsed with TBS buffer after 1 h incubation at RT and used immediately. Subsequent HS-AFM measurements were carried out on particles that had an average size of 550 ± 260 nm (n = 14) with 5 gel-holes being analyzed per particle on average. In some cases, Kap95 was added to the imaging buffer to achieve a final concentration of 1 µM Kap95.

#### Nup100 hydrogel permeation assay

To obtain 10 µl droplets containing micro-condensates 0.3 µl of 100 µM yNup100FG fragments labelled with fluorescein-5-maleimide (yNup100FG-FMAL) (in 2 M Guanidine-HCl, 100 mM Tris-HCl, pH 8) were diluted out in 9.7 µl of TBS and incubated at 25°C for 1 h. Kap95 was labeled with Alexa Fluor568 C5 Maleimide (Thermo Fischer) and purified on the spin column (Princeton separations) to remove excess dye. Then 5 µl of Kap95 was added to the 10 µl droplet containing yNup100FG-FMAL micro-condensates (final concentration ca. 10 µM) during transport and FRAP assays. BSA was labeled with Alexa Fluor647 C2 Maleimide (Thermo Fischer) and used as a non-specific reporter to test for passive permeability.

#### Fluorescence recovery after photobleaching

FRAP measurements were conducted using a point scanning confocal LSM880 Inverted microscope with Airyscan detector (Zeiss) built on a Zeiss Axio Observer stand (Intelligent Imaging Innovations GmbH). The system is equipped with a 1.4NA 63x plane apochromat objective (Plan-Apochromal 63x/1.4 Oil DIC M27), EMCCD camera (Evolve(R) 512, Photometrics) and a humidified climate control system at 25°C. A round 1 μm^2^ region within each condensate was chosen and bleached by application of three 10 ms pulses per pixel using solid state 488 nm laser (10 mW) for yNup100 FRAP or 555 nm laser (10 mW) for Kap95 FRAP, respectively. Sequential images (including 5 images before and 115 images after the bleaching event) were collected every 5 seconds by illuminating the sample with a 488 nm or with a 555 nm laser at 30% of its power (10 mW), respectively. Collected movies were analyzed using ImageJ, movies were checked for the oversaturated pixels (HiLo) and corrected for mobility of the condensates (stack registration). For each movie frame a time-stamp and fluorescent intensities of bleached area, whole condensate and background were exported and collected in an EXCEL file. These data were used to plot and analyze recovery curves using easyFRAP software (Markaki and Harz, 2017; Rapsomaniki et al., 2012).

#### In vivo localization assay of NupFG domain biomolecular condensates

Budding yeast *S. cerevisiae* strains used in this study are listed in the Key Resources Table. To assess the localization of the overexpressed NupFG domains, cells containing the pGAL-GFP-NupFG::His3 plasmids were grown at 30°C for 1 day on SD medium lacking histidine, containing 2% glucose (w/v), and then cultured 1 day on medium containing 2% raffinose (w/v) as carbon source. On the day of the experiment, NupFG expression was induced in exponentially growing cells with 1% (w/v) D-galactose for 1h. pACM021-GFP plasmid backbone, derived from pUG34, was used for the over-expression of the different NupFG domains. Selection of the FG domains of FG nucleoporins was based on (Yamada et al., 2010). For Nup153FG, the selection of the FG domain was based on (Kuiper et al., 2022). Details of the primers and plasmids used are listed in the Key Resources Table. To assess the localization of the different full-length FG nucleoporins, tagged with GFP at the C terminus, expressed under their endogenous promoters, cells were grown in 2% D-glucose SD medium lacking histidine.

To assess the localization of the overexpressed GFP-Nup100FG in the W303, Mlp1, ΔFG, and Nsp1FGx2 stains, cells containing the genome integrated TEFF-mCherry-WALP-HDEL::Leu2 (NE/ER marker) and the replicative plasmid pGAL-GFP-Nup100FG::Ura3 were grown in SD medium lacking leucine and uracil. For the ΔNsp1 p410-Nsp1FGx2:KanMX strain, medium was supplemented with G418 (200 μg/ml). Over-expression of Nup100FG was induced as explained previously (1% (w/v) D-galactose for 1h), before imaging.

For assessing the solubility of GFP-Nup100FG condensates, GFP-Nup100FG was over-expressed for 1h in the indicated strains, and cells were treated for 10 minutes with 10% 1,6-hexanediol (1,6HD), an aliphatic alcohol that dissolves liquid particles, before imaging.

Imaging was done on a DeltaVision Deconvolution Microscope (Applied Precision (GE), Issaquah, WA, USA), using InsightSSITM Solid State Illumination at 488 and 594 nm and an Olympus UPLS Apo 100× oil objective with 1.4 numerical aperture. Detection was done with a PCO-edge sCMOS camera (Photometrics, Tucson, AZ, USA). Image stacks (30 stacks of 0.2 µm) were deconvolved using softWoRx (Cytiva). Data were analyzed with open-source software ImageJ.

#### Cell lysis and western blot

Budding yeast cells overexpressing GFP-Nup100FG for 1h were collected by centrifugation, washed with PBS, and frozen in liquid nitrogen. Cell pellets were resuspended in HEPES buffer (50 mM HEPES, pH 7.5, 100 mM NaCl, 2.5 mM MgCl_2_, 10 mM DTT and 10% glycerol) supplemented with protease inhibitors (10 mM PMSF and cOmplete-EDTA protease inhibitor cocktail), and broken with glass bead using the fast-prep homogenizer. The lysate was clarified by centrifugation at 4,000 x rpm for 3 minutes. Total protein concentration of whole cell extracts was quantified using Pierce^TM^ BCA Protein Assay Kit.

To assess GFP-Nup100FG protein levels by western blot, equal amounts of whole cell lysates of the strains W303, Mlp1, ΔFG, and Nsp1FGx2 were diluted in 2X protein loading dye buffer and boiled for 15 min. Subsequently, samples were separated via SDS-PAGE (Stain free gels), transferred to PVDF membranes for 1h. Membranes were blocked with 2.5% BSA in PBS-T (0.1% Tween20), incubated overnight with primary antibody monoclonal mouse anti-GFP (1:2500) and secondary antibody anti-mouse (1:2500), and revealed with ECL using the Chemidoc imaging system (BioRad).

### QUANTIFICATION AND STATISTICAL ANALYSIS

#### Estimation of decay times from ACFs in HS-AFM data

To analyze the dynamic behavior over time at each radius, a kymograph was generated along a circle in a movie oriented radially around the central axis of each frame (**Fig S6**). The circle has radius *r* and any point along that circle has angle *θ*. For each frame in the movie, we assigned Euclidean *xy* coordinates to each pixel placing the origin at the exact center of the image, *c* = (0, 0). All pixels whose distance, *r_i_*, from *c* satisfy *r_i_* ϵ [*r* – *dr*,*r* + *dr*] (in which *d*r is defined less than half of the pixel size, and therefore the kymograph was taken at 1-pixel intervals in the radial direction) are sorted by *θ*_i_ and then sorted by *r_i_*. We repeated the process to generate kymographs from the maximum radius *r_n_* to the minimum radius of 0.5 pixel. The y- and x-axis in each kymograph denote angle and time, respectively. The analysis was carried out for HS-AFM movies recorded longer than 30 seconds (i.e., 200 or more frames recorded at 0.15 s per frame).

The ACFs were applied to all kymographs to quantify height changes, expressed as (Stanley et al., 2019):

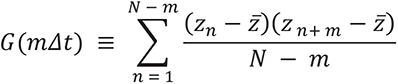

where m and n refer to the number of columns in the kymograph, N is the total number of columns, *z_n_* is the pixel intensity at the given frame and *z̄* is the mean value of *z_n_*. The longer m is, the lower number of data points are available to compute the ACF. Therefore, m was limited to N/2 or less for the ACF accuracy. Each ACF was normalized by the value at lag 0 and averaged over along the angle to generate an averaged single row. Each of the averaged rows was then compiled into a single heatmap. The y- and x-axis are radius and time-lag, respectively. The average ACF plot was then fitted with a single exponential decay model to estimate a decay time (Tau). To exclude possible intrinsic HS-AFM noise, the average ACF plot was fitted starting at lag 1 because the autocorrelation coefficient at lag 1 could be computed mostly from intrinsic noise. Furthermore, given the fact that autocorrelation coefficients become less reliable as the time-lag increases, only the coefficients above a threshold of 95% confidence interval were fitted (Christodoulou et al., 2021).

#### Assessment of Nup100FG localization and solubility to 1,6HD in cells

Unless otherwise stated three independent repeats of each experiment were performed. For assessing Nup100FG localization, condensates of fifty cells were analyzed for each replicate, using ImageJ. For assessing Nup100FG solubility to 10% 1,6HD, the number of condensates of cells untreated versus 10% 1,6HD treated was quantified. Graphs and statistical analyses were generated using Prism software (GraphPad). Band intensities following immunoblotting and detection using chemiluminescence reagents (ECL) were quantified with ImageJ. Individual results from all three repeats are shown, together with the means and standard deviations. For significance analysis, we used unpaired Student’s t tests. ∗ is refers to p < 0.05, ∗∗ is refers to p < 0.01, and ∗∗∗ is refers to p < 0.001. Further details about the number of samples can be found in the corresponding figure legends.

## Notes

### Competing Interest Statement

The authors have declared no competing interest.

### Summary of Updates

Fig. 3 and Fig. S10 now include additional experimental replicates. Updated author list.

