## Supplemental Figures, Tables and Movie legends for "Dynamic molecular mechanism of the nuclear pore complex permeability barrier"

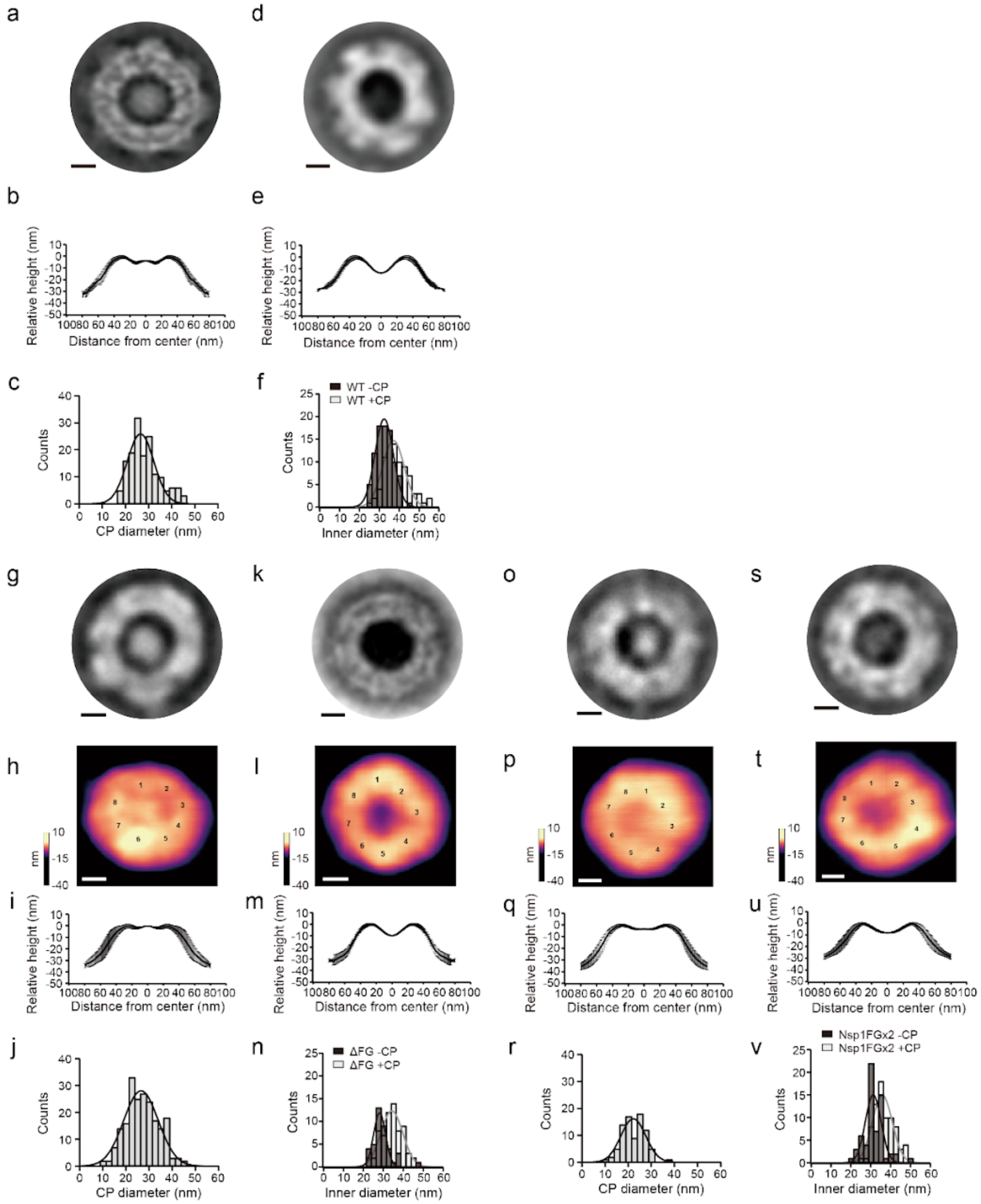


**Figure S1. WT and mutant NPCs retain their overall structure. a-f,** Average negative stain TEM images of +CP (**a**; n = 172) and of -CP (**d**; n = 76) WT NPCs. Average cross-sectional profiles of +CP (**b**; n = 10) and -CP (**e**; n = 19) NPCs that correspond to the HS-AFM images shown in Figs. 1b and 1d. Histograms of CP diameters (**c**, mean value is 29.4 ± 7.7 nm; 52 NPCs) and inner diameters **(f)** for +CP WT NPCs (gray, mean value is 39.7 ± 6.1 nm; n = 21) and -CP WT NPCs (black, mean value is 33.8 ± 4.1 nm; n =

22) measured by HS-AFM. **g-n,** Average negative stain TEM and HS-AFM images of +CP ΔFG NPCs (**g**; n = 131 for TEM and **h**; n = 8 for HS-AFM) and -CP ΔFG NPCs (**k**; n = 184 for TEM. **l**; n = 8 for HS-AFM). Average cross-sectional profiles of +CP ΔFG NPCs (**i**; n = 8) and -CP ΔFG NPCs (**m**; n = 8) measured by HS-AFM. Histograms of CP diameters (**j**, mean value is 26.5 ± 10.5 nm; n = 69) and inner diameters **(n)** for +CP ΔFG NPCs (gray, mean value is 34.2 ± 7.8 nm; n = 17) and -CP ΔFG NPCs (black, mean value is 28.1 ± 4.2 nm; n = 10) measured by HS-AFM. **o-v,** Average negative stain TEM and HS-AFM images of +CP Nsp1FGx2 NPCs (**o**; n = 29 for TEM. **p**; n = 6 for HS-AFM) and -CP Nsp1FGx2 NPCs (**s**; n = 62 for TEM. **t**; n = 7 for HS-AFM). Average cross-sectional profiles of +CP Nsp1FGx2 NPCs (**q**; n = 6) and -CP Nsp1FGx2 NPCs (**u**; n = 7) measured by HS-AFM. Histograms of CP diameters (**r**; mean value is 22.2 ± 7.5 nm; n = 28) and inner diameters **(v)** for +CP Nsp1FGx2 NPCs (gray, mean value is 35.8 ± 7.1 nm; n = 19) and -CP Nsp1FGx2 NPCs (black, mean value is 31.2 ± 5.6 nm; n = 16) measured by HS-AFM. Numbers on the HS-AFM average images indicate eightfold rotational symmetry. Scale bars, 20nm.

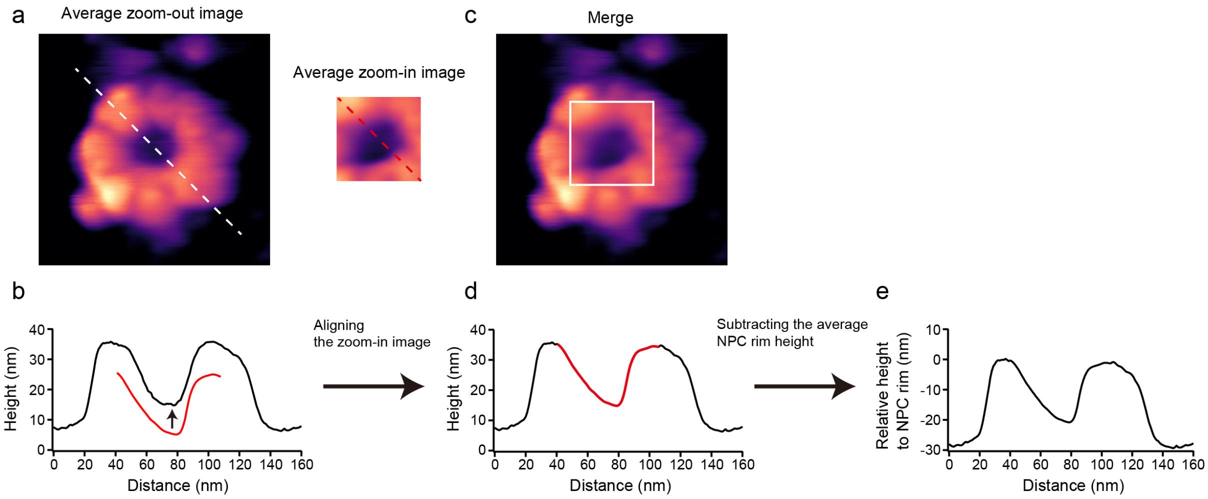


**Figure S2. Alignment of zoom-in to zoom-out images. a,** Average zoom-out and zoom-in images of an NPC channel. **b,** The cross-sectional profile taken along the white and red dashed lines in **(a)**. **c-d,** The zoom-in image is aligned with the zoom-out image in the x- and y-directions **(c)** and the z-direction **(d)**. **e,** From the zoomed-out image, the average maximum height (termed NPC rim) is calculated from the 8 spokes of each NPC and is subtracted from the zoom-in image height. In this way, the relative height of features in a zoom-in image is referenced to a zero-height value that is assigned to the NPC rim.

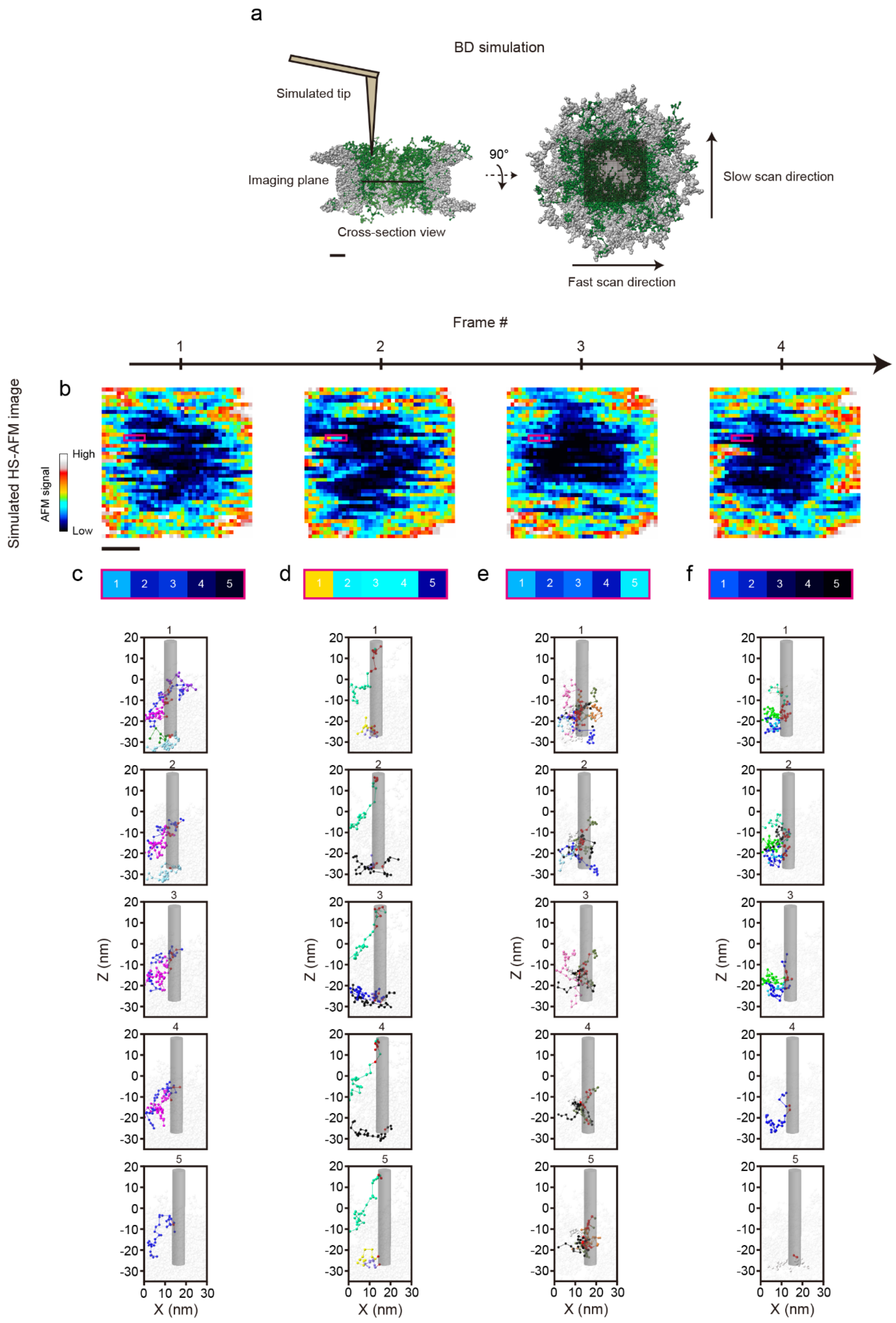
a
BD simulation
Simulated tip
Imaging plane
Cross-section view
90°
Slow scan direction
Fast scan direction
Frame #
1
2
3
4
b
Simulated HS-AFM image
AFM signal
High
Low
c
d
e
f
Z (nm)
X (nm)

**Figure S3. HS-AFM downsamples FG domain dynamics. a,** HS-AFM simulation using a tip with 3 nm-radius that scans over a Brownian Dynamics (BD) model of the isolated yeast NPC. Scaffold, gray; FG domains, green. The total run time of the HS-AFM simulation is 72 µs. Right panel shows the top view of the NPC model highlighting the 40 x 40 pixel scan area of the HS-AFM simulation. **b,** Successive HS-AFM images of the BD model obtained by a simulated raster scan of the HS-AFM tip at a rate of 16 us per frame. The time required to capture one pixel of data in the HS-AFM image (i.e., "scan speed") is 0.01 µs per pixel, which is equivalent to the duration of one BD "snapshot". The scan direction is from left to right starting from the bottom left corner. Note that horizontal stripes dominate images. Scale bar, 10 nm. **c-f,** Pixel values within the stripe bounded by the magenta box in Frames 1, 2, 3 and 4, are derived from contact between a 3 nm-radius HS-AFM tip (gray cylinder) and the FG domain "beads" beneath it (red). Otherwise, each individual FG domain is assigned a unique colour across all snapshots. All other beads are transparent. The difference in stripe behaviour between frames originates from the presence of different FG domain beads that contact the HS-AFM tip in the time that has elapsed from Frame 1 to Frame 4.

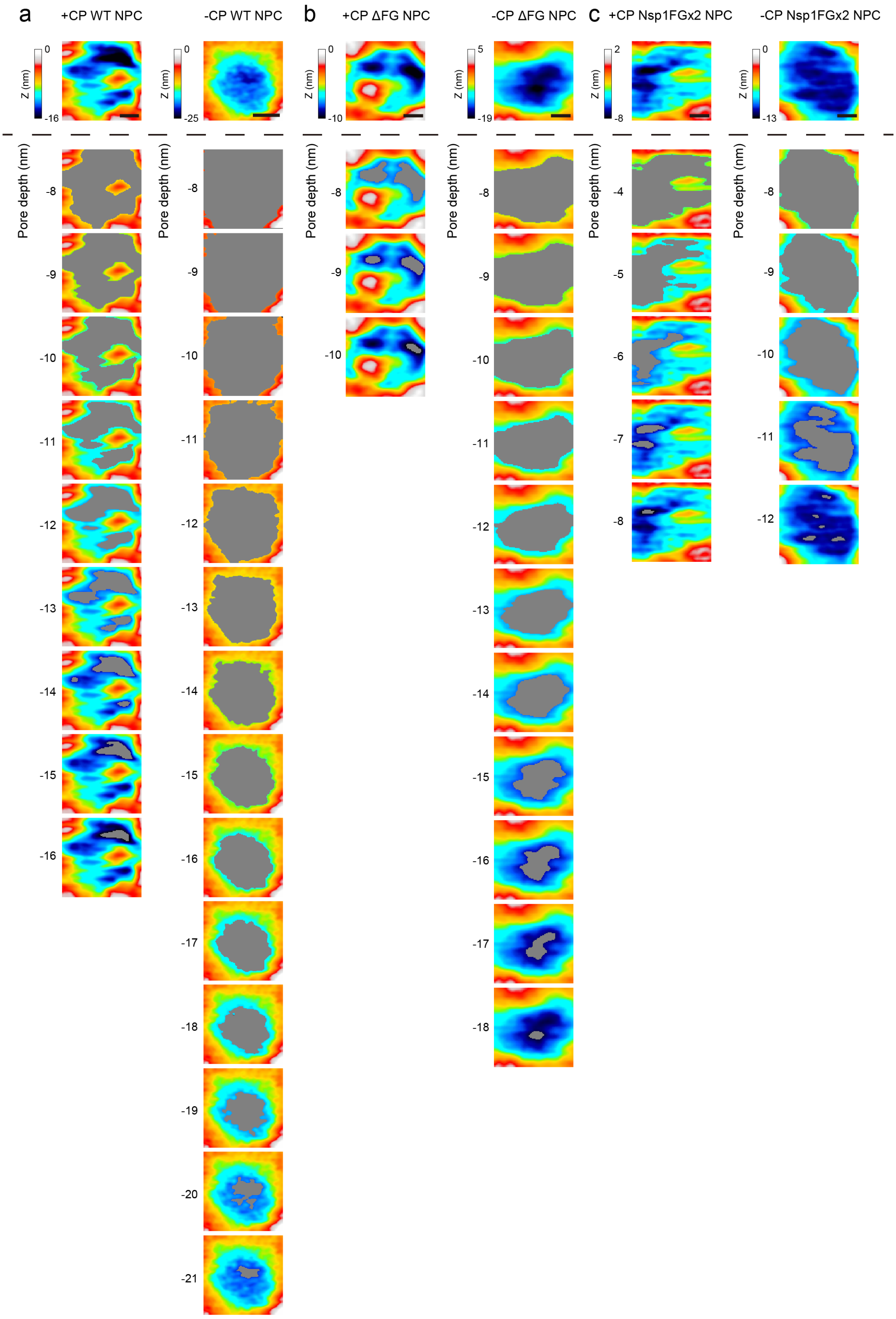


**Figure S4. Visualizing voids at 1 nm increments in pore depth. a-c,** Representative HS-AFM images of WT **(a)**, ΔFG **(b)** and Nsp1FGx2 **(c)** NPCs as shown in Figures 1, 2 and 3, respectively. Voids (gray) are shown at 1-nm increments in pore depth. Top row: Corresponding full scale topographical images. Scale bars, 10 nm.

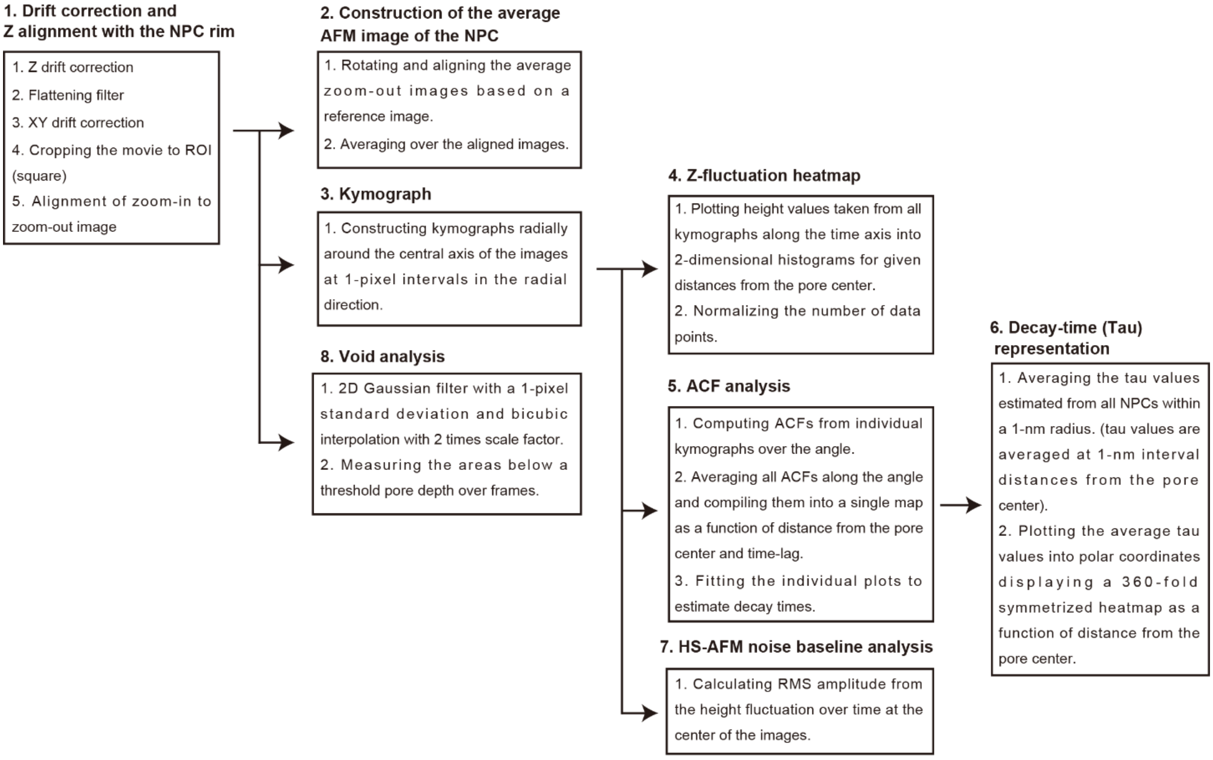


**Figure S5. HS-AFM analysis workflow.** See Methods for more details.

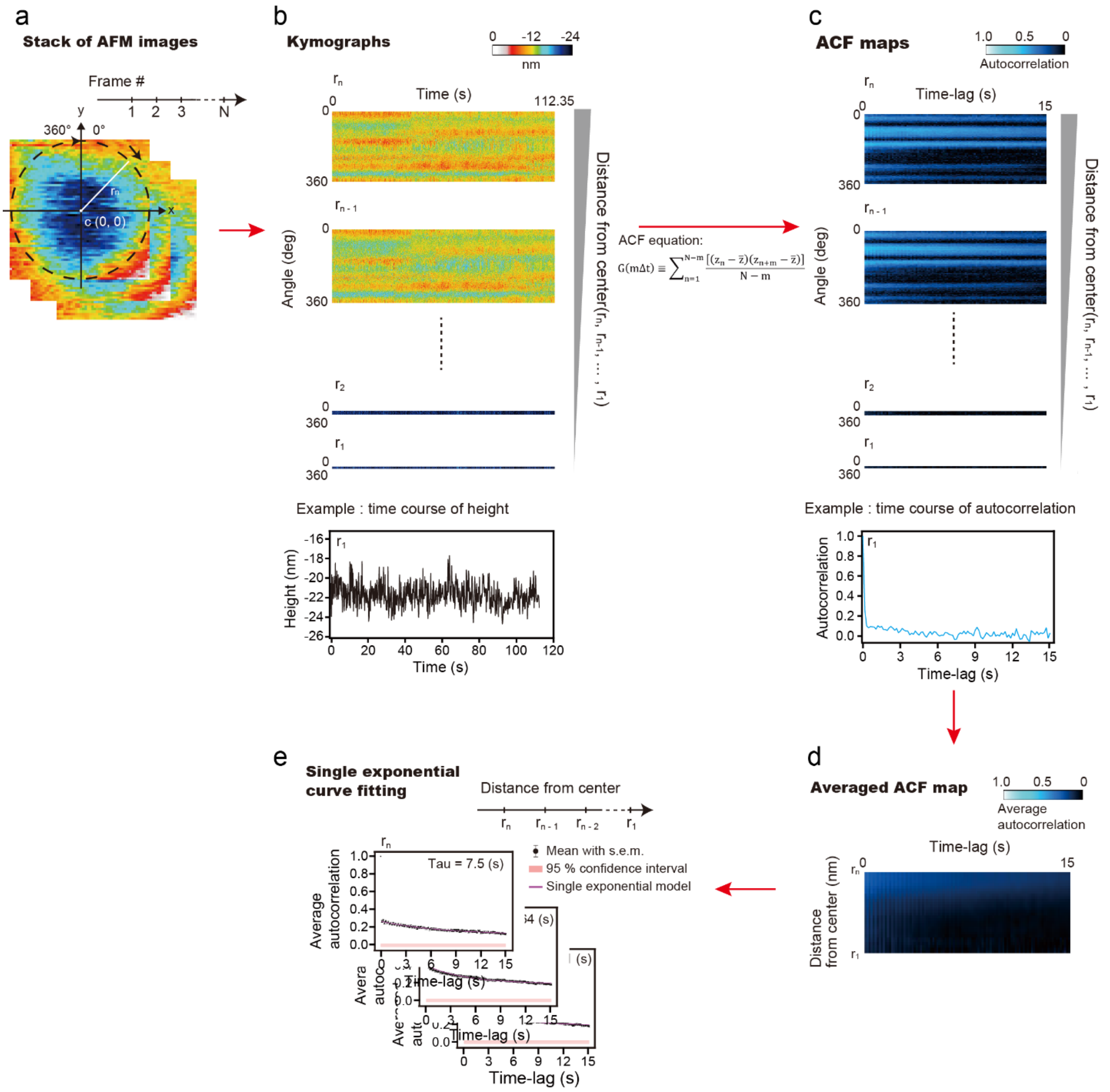


**Figure S6. ACF analysis workflow. a,** Stack of HS-AFM images with xy coordinates assigned to each pixel placing the origin at the centre of the image. Kymographs are taken clockwise along the circumference from 0 to 360 degrees (θ) at a distance of radius $r_n$ (indicated by white line) away from the origin c = (0, 0). **b,** Kymographs taken from the maximum radius $r_n$ to the minimum radius $r_1$. The x- and y-axes are time and angle, respectively. Bottom, example of a height trace taken from the kymograph $r_1$. **c,** Normalized ACF maps computed from each of the kymographs. Bottom, example of an autocorrelation coefficient trace taken from the ACF map $r_1$. **d,** Each of the ACF maps in **(c)** were averaged over along the angle, and then compiled into a single map. The x- and y-axes are time-lag and radius, respectively. **e,** The ACF plot at each radius was fitted with a single exponential decay model to estimate a decay time (Tau).

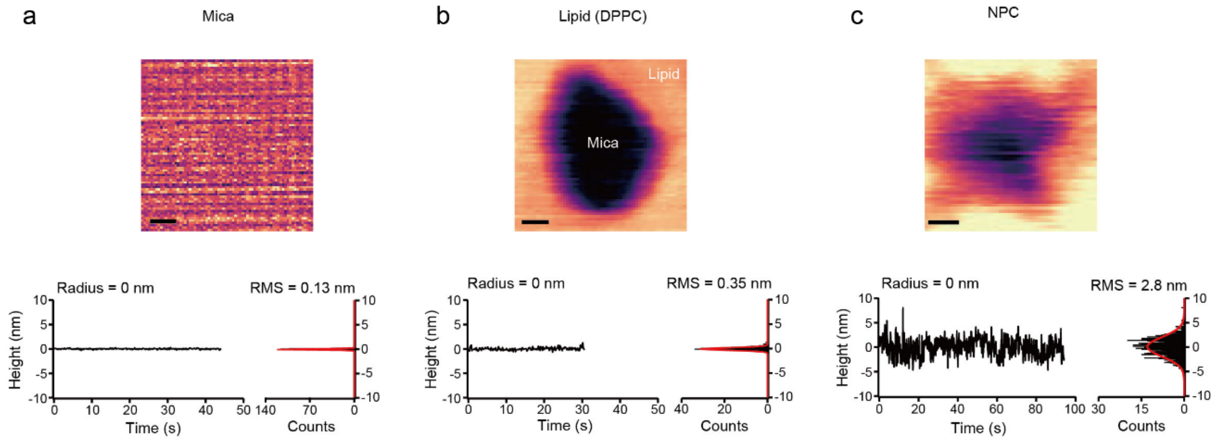


**Figure S7. HS-AFM noise baseline analysis. a-c,** HS-AFM images of a mica surface **(a)**, a lipid membrane (100% DPPC) with a defect **(b)** and an NPC channel **(c)**. Each height trace over time on the bottom left corner was taken at the centre of the image. Histograms with a gaussian distribution fit on the right of the trace for mica, lipid and NPC give RMS values of 0.13 nm, 0.35 nm and 2.8 nm, respectively. Scale bars, 10 nm.

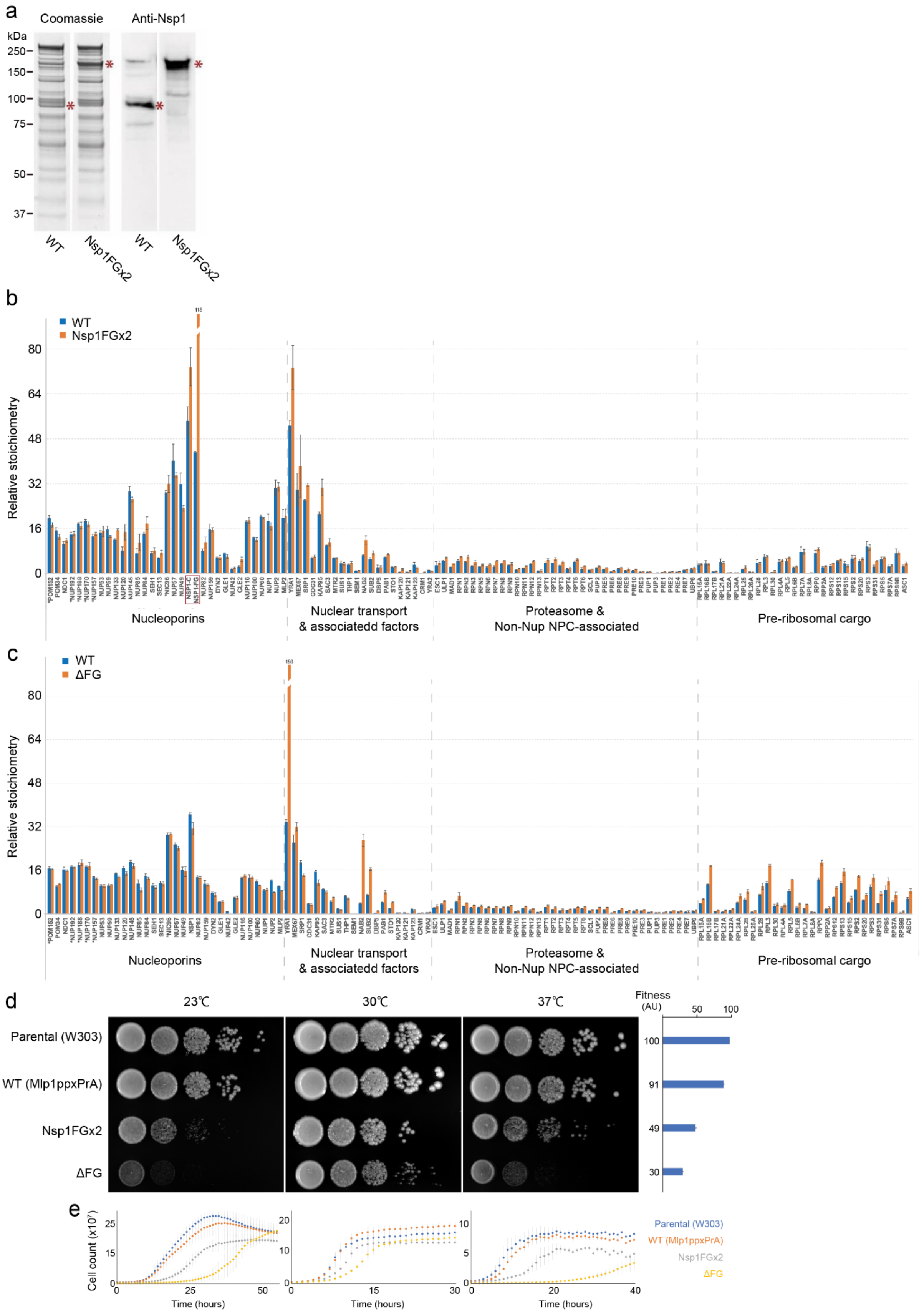


**Figure S8. Characterization and fitness of Nsp1FGx2 mutant cells compared with WT and ΔFG strains. a,** Expression and molecular weight of the Nsp1FGx2 mutant protein in affinity-purified NPCs shown by Coomassie staining (left) and western-blot using an anti-Nsp1 antibody (right). Asterisks indicate the position of the wild type Nsp1 and mutant Nsp1FGx2 proteins. **b-c,** Comparative relative stoichiometry of Nups and main associated proteins within affinity-purified NPCs (Kim et al., 2018) between wild type (blue bars) and the Nsp1FGx2 (b) and ΔFG NPC (c) mutants (orange bars), as determined by label-free MS quantification (at least 5 peptides per protein). Proteins are grouped by functional categories or membership of discrete macromolecular assemblies (in some cases only a selection of components are shown). The bars for Nsp1 and Yra1 were shortened for presentation purposes, with their actual value indicated above. In the Nsp1FGx2 plot, two values are provided for Nsp1, indicating the values obtained when only analyzing peptides originated from the C-terminal anchor site of the protein (NSP1-C), or peptides originated only from the FG region of the protein (NSP1-FG), reflecting the levels of incorporated protein and FG repeats within the NPCs respectively. The handle used for the affinity purification (Mlp1) is not shown. Biological replicas, n = 3. Error bars = standard deviation. **d,** Spot growth tests at different temperatures (23°C, 30°C and 37°C) for the parental (w303) and mutant strains Mlp1ppxPrA, Nsp1FGx2 Mlp1ppxPRA and ΔFG Mlp1ppxPrA strains. Serial 10-fold dilutions of cells were spotted on YEPD plates and grown at the indicated temperatures for 1–3 d. Each growing phenotype was quantified by semiquantitative methods (see STAR Methods), and the obtained value (in arbitrary units [AU]) is shown on the right of each column. Plotted fitness value (mean ± s.d.) for each measurement is shown on the right. **e,** Growth curves for the same strains and temperatures as in (d) measured in 96-well plates for the times indicated. $OD_{600}$ measurements were transformed to cells/ml (x$10^7$) as indicated in the STAR Methods section. Biological replicas, n = 3-7. Error bars = standard deviation.

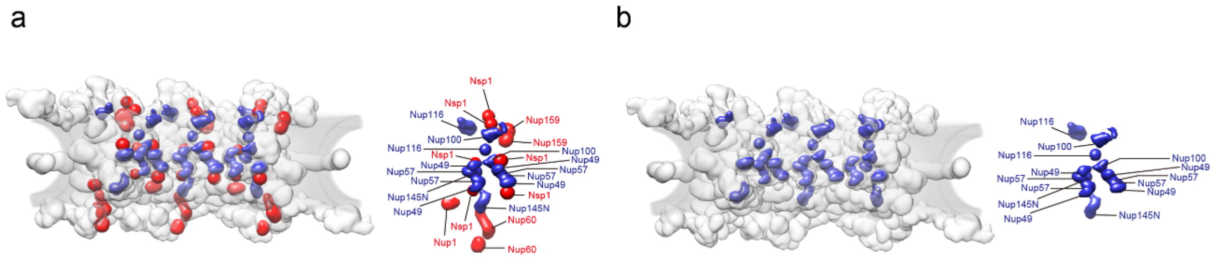


**Figure S9. FG domain anchor sites on the NPC scaffold. a-b,** Side view of three spokes showing the FG domain anchor sites for FxFG (red) and GLFG (blue) Nups in WT **(a)** and ΔFG NPCs **(b)**. Each view is accompanied by a single spoke with specific FG domain labelled.

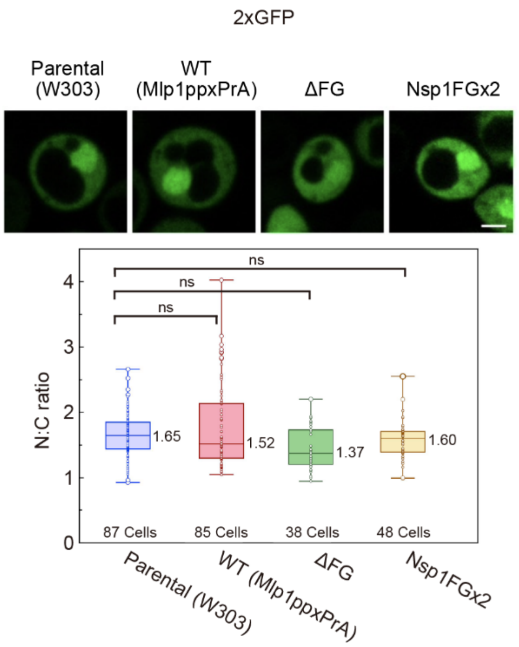


**Figure S10. Localization of 2xGFP in yeast cells.** The steady-state distribution (N:C ratio) of 2xGFP in Parental, WT, ΔFG and Nsp1FGx2 cells (no. of replicates = 3). Number of cells analyzed, median values, first and third quartiles are indicated in the box plots. Scale bar, 2 µm.

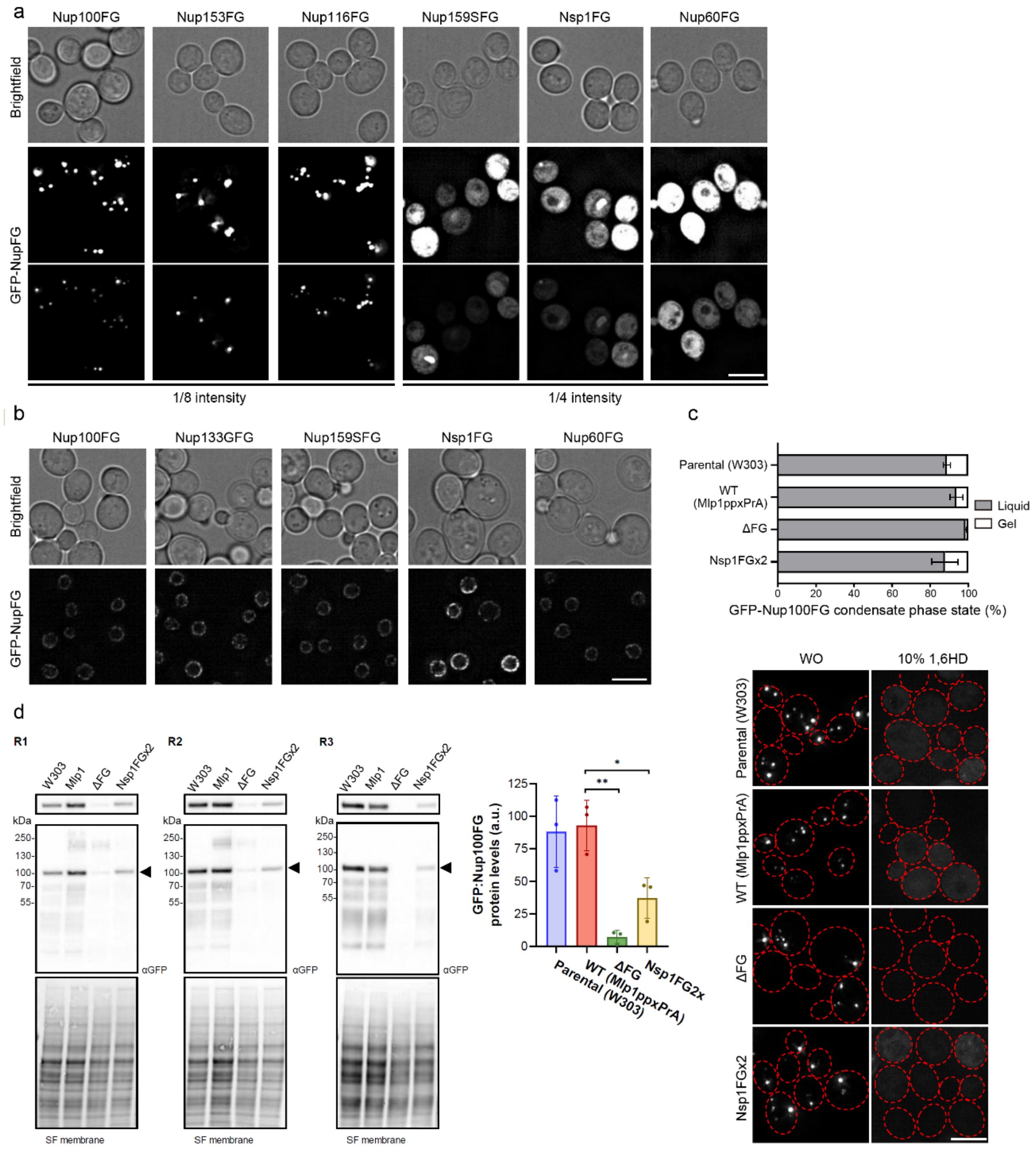


**Figure S11. Characterization of *in vivo* FG Nup condensates. a**, Cellular localization of the different FG Nup domains tagged with GFP, over-expressed for 1h in wt strain. Maximal projection of 4 z-stacks, 0,2 µm interval. Scale bar, 5 µm. The bottom images are as in **Fig. 5a** and have 8-fold (Nup100FG, Nup116FG, and Nup153FG) and 4-fold (Nup159SFG, Nsp1FG, and Nup60FG) lower intensity compared to the top panels. Top panels serve to assess if a punctate NPC signal becomes visible under the same imaging conditions used in **S11b** for imaging full length Nups. **b**, Localization of different full-length FG Nups, tagged with GFP at the C-term, expressed under their endogenous promoter. Maximal projection of 4 z-stacks, 0,2 µm interval. Scale bar, 5 µm. **c,** GFP-Nup100FG was over-expressed for 1h in the indicated strains and imaged without (WO) and after 10 min 10% 1,6-hexanediol (1,6HD) treatment. Graph shows the percentage of condensates that are in a liquid or gel phase state (based on their dissolution by the aliphatic alcohol 10% 1,6HD (no. of replicates = 3; mean ± s.d.). Cells analyzed: 65-300. **d,** Western blot analysis of GFP-Nup100FG over-expressed for 1h in the indicated strains. Bar plot represents the quantification of GFP-Nup100FG protein levels relative to protein loading (stain free staining) (no. of replicates = 3; mean ± s.d.). SF = stain free. Primary ab: mouse anti-GFP (1:2500); secondary ab: anti-mouse (1:2500). Arrows point at GFP-Nup100FG.

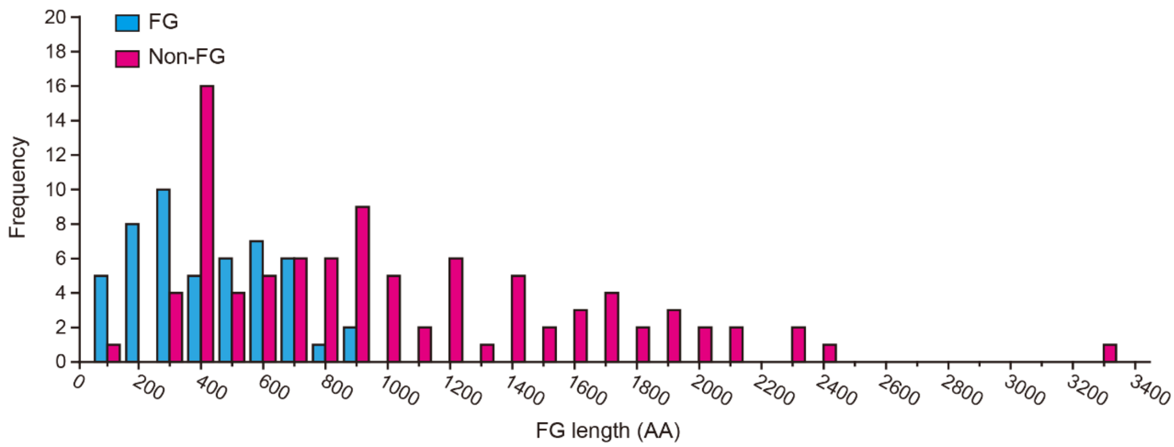


**Figure S12. Amino acid lengths for intrinsically disordered FG regions and structured Nups**. Histograms of amino acid (AA) lengths of FG regions (blue) and non-FG Nups (red) from 4 different species *(S. pombe*, *S. cerevisiae*, *H. sapiens* and *A. thaliana*). Several lengths of the FG regions are restrained while the lengths of the non-FG Nups are not constrained.

### Supplemental tables

**Table S1. Properties of *S. cerevisiae* FG Nups**

| S. cerevisiae | AAs in disordered domain[a] | Predicted hydrodynamic diameter (nm)[a] | Cohesiveness[a] | Copy number[b] | Number of FG repeats[c] |
|---|---|---|---|---|---|
| Nup159 | 387-1071 | 13.4 | Non-cohesive | 16 | 25 |
| Nup100 | 1-610 | 9.8 | Cohesive | 16 | 44 |
| Nup100 | 611-800 | 7.4 | Non-cohesive | | |
| Nup116 | 172-764 | 9.8 | Cohesive | 16 | 42 |
| Nup116 | 765-960 | 7.6 | Non-cohesive | | |
| Nup42 | 1-382 | 8.2 | Cohesive | 8 | 18 |
| Nup49 | 1-251 | 5.0 | Cohesive | 32 | 16 |
| Nsp1 | 1-186 | 6.2 | Cohesive | 48 | 33 |
| Nsp1 | 187-617 | 12.4 | Non-cohesive | | |
| Nup57 | 1-255 | 6.8 | Cohesive | 32 | 16 |
| Nup145N | 1-242 | 5.0 | Cohesive | 16 | 11 |
| Nup145N | 243-433 | 6.4 | Non-cohesive | | |
| Nup1 | 220-797 | 13.6 | Non-cohesive | 8 | 17 |
| Nup1 | 798-1076 | 7.2 | Cohesive | | |
| Nup2 | 160-800 | 12.6 | Non-cohesive | 16 | 12 |
| Nup60 | 389-539 | 6.6 | Non-cohesive | 16 | 4 |

[a]Based on Yamada et al., 2010.
[b]Based on Kim et al., 2018 and Huang et al., 2020.
[c]Number of FG repeats is taken from Uniprot (https://www.uniprot.org/)
AAs = amino acids

**Table S2. FG repeat and FG domain concentrations in yeast WT, ΔFG and Nsp1FGx2 NPCs and *X. laevis* NPCs**

| | Volume | FG repeat | | | FG domain | | |
|---|---|---|---|---|---|---|---|
| | $nm^3$ | $/nm^3$ | mg/ml | mM | $/nm^3$ | mg/ml | mM |
| **Isolated WT yNPC[a]** | ~157,963[c] | ~0.032 | ~13 | ~53 | ~0.572 | ~96 | ~2.3 |
| **Isolated ΔFG yNPC[a]** | ~157,963[c] | ~0.016 | ~6 | ~27 | ~0.237 | ~40 | ~1.1 |
| **Isolated Nsp1FGx2 yNPC[a]** | ~157,963[c] | ~0.041 | ~16 | ~69 | ~0.740 | ~124 | ~2.3 |
| **In situ WT yNPC[a]** | ~194,190[c] | ~0.026 | ~10 | ~43 | ~0.465 | ~78 | ~1.9 |
| ***Xenopus laevis* NPC[b]** | ~280,236[d] | ~0.019 | ~7 | ~31 | ~0.370 | ~61 | ~1.6 |

[a]Numbers of FG repeats are taken from Uniprot (https://www.uniprot.org/). FG domain copy numbers are based on Kim et al., 2018 and Huang et al., 2020.

[b]Numbers of FG repeats are taken from Uniprot. FG domain copy numbers are based on Labokha et al., 2013 and Zhu et al., 2022.

[c]Isolated yNPC and in situ yNPC volumes are calculated from Kim et al., 2018 and Akey et al., 2022, respectively.

[d]*Xenopus laevis* NPC volume is calculated from Eibauer et al., 2015.

#### Supplementary Movie Legends:

**Supplementary Movie 1.** HS-AFM movie of a +CP WT NPC recorded within 69 x 69 $nm^2$ with 80 x 80 pixels at 150 ms per frame (6.7 fps). The movie was cropped and processed using a 2D Gaussian filter with a 1-pixel standard deviation followed by a 10x scaling factor with bicubic interpolation. Magenta arrow indicates the CP. The playback speed is in real time. Scale bar, 10 nm.

**Supplementary Movie 2.** BD simulation of FG Nup dynamics in the NPC. Scaffold, gray; FG Nups, green. The total run time of the simulation is 72 µs (7,200 frames in total). The movie is played for the first 10 µs (i.e., 1,000 frames). The playback speed is 25 fps. Scale bar, 20 nm.

**Supplementary Movie 3.** A simulated HS-AFM movie of a BD simulated NPC. Each image was obtained by raster scanning a simulated 3-nm radius HS-AFM tip at a rate of 16 µs per frame. The simulated HS-AFM scan was initiated at the midplane of the NPC within an area of 39 x 39 $nm^2$ with 40 x 40 pixels. Each pixel was simulated from one BD snapshot. The playback speed is 2 fps. Scale bar, 10 nm.

**Supplementary Movie 4.** HS-AFM movie of a -CP WT NPC recorded within 39 x 39 $nm^2$ with 80 x 80 pixels at 150 ms per frame (6.7 fps). The movie was cropped and processed using a 2D Gaussian filter with a 1-pixel standard deviation followed by a 10x scaling factor with bicubic interpolation. The playback speed is in real time. Scale bar, 5 nm.

**Supplementary Movie 5.** HS-AFM movie of a +CP ΔFG NPC recorded within 79 x 79 $nm^2$ with 80 x 80 pixels at 150 ms per frame (6.7 fps). The movie was cropped and processed using a 2D Gaussian filter with a 1-pixel standard deviation followed by a 10x scaling factor with bicubic interpolation. Magenta arrow indicates the CP. The playback speed is in real time. Scale bar, 10 nm.

**Supplementary Movie 6.** HS-AFM movie of a -CP ΔFG NPC recorded within 79 x 79 $nm^2$ with 80 x 80 pixels at 150 ms per frame (6.7 fps). The movie was cropped and processed using a 2D Gaussian filter with a 1-pixel standard deviation followed by a 10x scaling factor with bicubic interpolation. The playback speed is in real time. Scale bar, 10 nm.

**Supplementary Movie 7.** HS-AFM movie of a +CP Nsp1FGx2 NPC recorded within 79 x 79 $nm^2$ with 80 x 80 pixels at 150 ms per frame (6.7 fps). The movie was cropped and processed using a 2D Gaussian filter with a 1-pixel standard deviation followed by a 10x scaling factor with bicubic interpolation. Magenta arrow indicates the CP. The playback speed is in real time. Scale bar, 10 nm.

**Supplementary Movie 8.** HS-AFM movie of a -CP Nsp1FGx2 NPC recorded within 79 x 79 $nm^2$ with 80 x 80 pixels at 150 ms per frame (6.7 fps). The movie was cropped and processed using a 2D Gaussian filter with a 1-pixel standard deviation followed by a 10x scaling factor with bicubic interpolation. The playback speed is in real time. Scale bar, 10 nm.
